# Exploring Abeta42 Monomer Diffusion Dynamics on Fibril Surfaces through Molecular Simulations

**DOI:** 10.1101/2024.02.29.582685

**Authors:** Yuan-Wei Ma, Guan-Fang Wang, Hong-Yi Chen, Min-Yeh Tsai

**Affiliations:** Institute of Bioinformatics and Structural Biology, National Tsing-Hua University, Hsinchu, 300044, Taiwan; Department of Chemistry and Biochemistry, National Chung Cheng University, Minhsiung, Chiayi 621301, Taiwan; Center for Nano Bio-Detection, National Chung Cheng University, Minhsiung, Chiayi 621301, Taiwan; Division of Physics, National Center for Theoretical Sciences, National Taiwan University, Taipei 106319, Taiwan

## Abstract

This study provides critical insights into the role of surface-mediated secondary processes in Alzheimer’s disease, particularly regarding the aggregation of Abeta42 peptides. Employing coarse-grained molecular dynamics simulations, we focus on elucidating the molecular intricacies of these secondary processes beyond primary nucleation. Central to our investigation is the analysis of a freely diffusing Abeta42 monomer on pre-formed fibril structures. We conduct detailed calculations of the monomer’s diffusion coefficient on fibril surfaces (as a one-dimensional case), along with various monomer orientations. Our findings reveal a strong and consistent correlation between the monomer’s diffusion coefficient and its orientation on the surface. Further analysis differentiates the effects of parallel and perpendicular alignments with respect to the fibril axis. Additionally, we explore how different fibril surfaces influencèmonomer dynamics by comparing the C-terminal and N-terminal surfaces. We find that the monomer exhibits lower diffusion coefficients on the N-terminal surface. Differences in surface roughness (S_R_), quantified using root-mean-square distances, significantly affect monomer dynamics, thereby influencing the secondary aggregation process. Importanly, this study underscores that fibril twisting acts as a regulatory niche, selectively influencing these orientations and their diffusion properties necessary for facilitating fibril growth within biologically relevant time scales. This discovery opens new avenues for targeted therapeutic strategies aimed at manipulating fibril dynamics to mitigate the progression of Alzheimer’s disease.

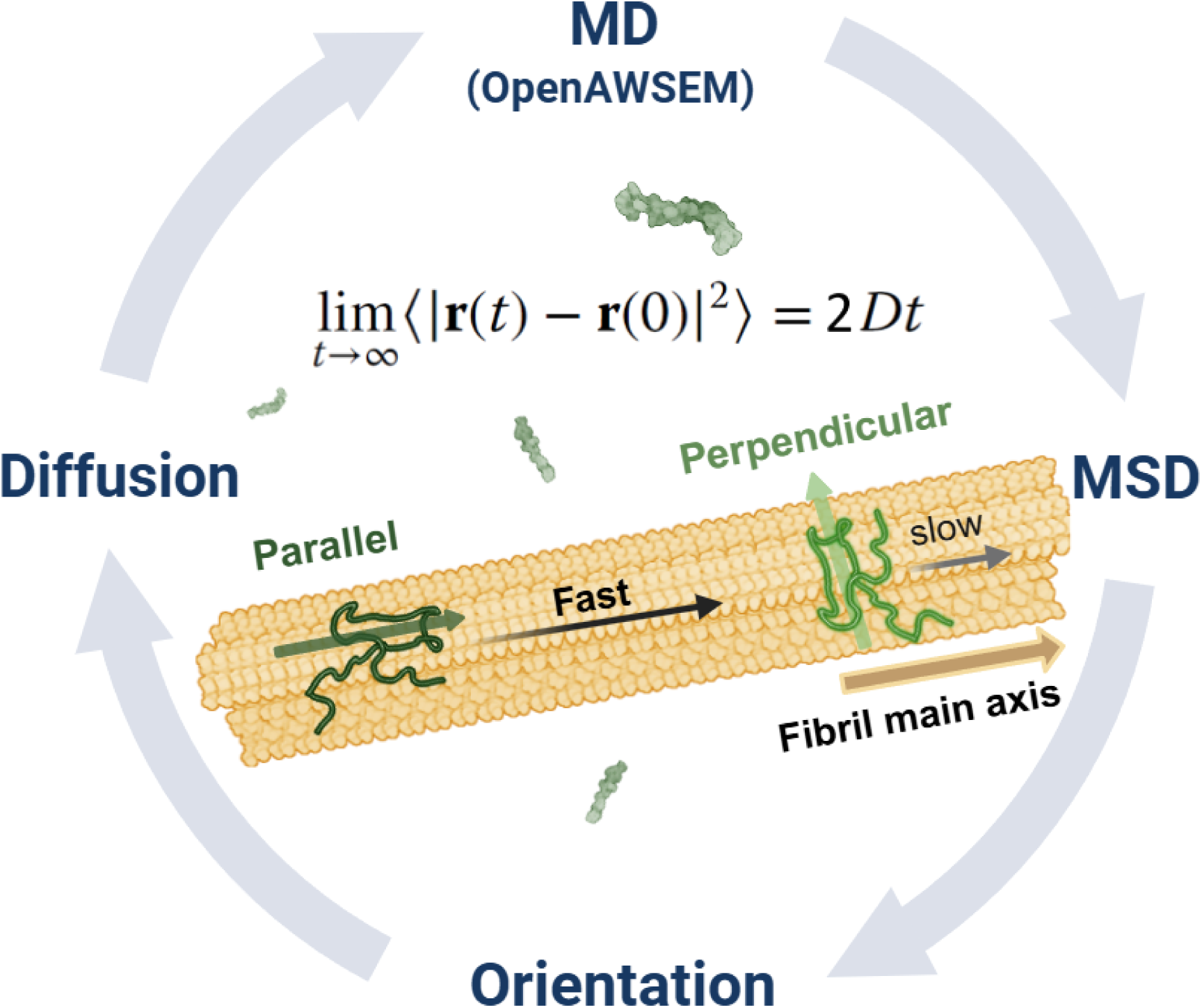

## 1 INTRODUCTION

Protein aggregation plays a pivotal role in diseases such as Alzheimer’s, especially in the context of the Amyloid Hypothesis.^1^ Understanding this process, which includes nucleation, fibril growth, and breakage, is crucial for developing potential therapeutic interventions.^2^ Despite this, the treatment landscape for Alzheimer’s faces significant challenges. This is evident in the case of FDA-approved drugs like Aducanumab (Aduhelm), which are limited by usage constraints and safety issues.^3,4^

Recent studies have put forward the ‘oligomer hypothesis,’ which posits that while amyloid fibers might represent stable thermodynamic end-products, the toxic elements predominantly emerge from oligomers formed during protein aggregation^5–7^ (see a recent comprehensive review^8^). Understanding of primary nucleation mechanisms in amyloid fibers is robust;^9–11^ however, the dynamic complexity of protein fibril growth, particularly at lower concentrations (less than 10 µM), continues to present challenges.^12^ Insights from examining the morphologies and shapes of Abeta oligomers have deepened our understanding of how these oligomers, especially those with lateral branching, are formed.^13^ Nonetheless, secondary processes such as filament breakage and secondary nucleation introduce additional complexity, substantially affecting the progression of neurodegenerative diseases.^14,15^

Fibril growth and secondary processes are intricately linked to interactions at fibril surfaces and subsequent catalytic properties. Despite pioneering experiments elucidating surface-mediated nucleation (for example, sickle hemoglobin^16,17^ and amyloid proteins^15,18^), a significant knowledge gap persists in understanding the molecular mechanisms underpinning these processes. These processes are crucial as they significantly influence the formation of toxic oligomers,^19–21^ reflect reactive surface binding sites,^20,22^ and modulate the macroscopic ‘lag time’.^19^ This gap is particularly evident in the limited experimental data available for visualizing these interactions in real-time or at atomic resolution, as emphasized in recent reviews.^23^ Interestingly, Xu et al. employed time-resolved in situ scanning probe microscopy, revealing that the monomer’s association rate to the fibril end is significantly below the diffusion limit, indicating a high free energy barrier for new monomer incorporation into fibrils.^12^ The finding suggests that other rate-limiting processes, potentially involving surface-mediated interactions, could be influencing the monomer binding dynamics. Computational simulations can bridge this gap by providing a detailed understanding of these molecular interactions, which are challenging to capture directly through experimental methods.^23,24^ Full-atom simulations have shed light on early structural changes in aggregation, particularly hydrophobic and electrostatic forces influencing peptide adsorption on surfaces.^25,26^ Coarse-grained models better explore longer timescales and biological contexts, revealing multiple aggregation pathways influenced by peptide-surface interactions and rigidity.^27–31^ These computational insights have proven invaluable in identifying key features of protein aggregation that are difficult to observe experimentally, especially when considering the scale and complexity of biological systems.

Studies emphasize protein monomer mobility and lateral diffusion on surfaces in surface-mediated protein aggregation.^32–35^ A crucial aspect of this is the intricate balance in the energy of protein monomer-surface interactions, which lies within a very narrow range and fulfills the energy requirement for secondary nucleation.^36^ However, our understanding of monomer conformations and their diffusion behavior at the microdynamic level is still incomplete. For instance, the relationship between a protein’s conformation and its diffusion dynamics on fibril surfaces remains an intriguing question. Gaining a comprehensive insight into these aspects is essential for effectively controlling monomer dynamics on surfaces.

Understanding the dynamics of protein monomers on fibril surfaces is not just crucial for surface-mediated aggregation, but also connects to broader principles in physical science. This leads us to consider the fundamental role of diffusion in natural processes, a concept central to the fluctuation-dissipation theory.^37^ Diffusion, a key driver of atomic and molecular Brownian motion, manifests across various states of matter, including gases, liquids, solids, and glasses. Within biological contexts, such as in protein behavior, we observe significant variability in diffusion rates contingent on the environment. For example, protein diffusion is markedly slower in the crowded cytoplasm compared to dilute solutions (10 µm²/s vs. 100 µm²/s).^38^ Specifically, when proteins are attached to fibril surfaces, advanced techniques like Total Internal Reflection Fluorescence Microscopy (TIRFM) and Atomic Fluorescence Microscopy (AFM) have been instrumental in estimating diffusion coefficients, which range between 0.5-2.4 µm²/s.^34^

Our study therefore focuses on the ‘Oligomer Cascade Hypothesis,’ which asserts that the mechanisms of seeded polymerization and surface-catalyzed nucleation are crucial for the formation and neurotoxic effects of soluble amyloid oligomers, key agents in the progression of neurodegenerative diseases.^39,40^ These processes suggest that secondary processes on fibril surfaces depend on the dynamics and structure of newly attached protein monomers. Heterogeneous fibril surfaces, including assembly defects and specific amino acid binding sites, facilitate monomer diffusion, sliding, collisions, secondary nucleation, fibril elongation, and breakage, possibly leading to toxic aggregates. To explore this, we employ the openAWSEM simulation platform,^41^ known for its accuracy in physicochemical information, biological self-assembly, and fibril aggregation studies.^42–45^ openAWSEM excels in biophysical mesoscale kinetic analysis, capturing relevant time scales and molecular details, making it ideal for investigating the mesoscopic diffusion dynamics of amyloid-like peptides on fibril surfaces, a pioneering endeavor in this field.

By leveraging extensive simulations guided by the energy landscape theory,^46,47^ the goal is to unravel the role of surface interactions in governing fibril growth, potentially offering innovative insights for neurodegenerative disease treatment and management. This research aims to enrich theoretical understanding and guide experimental approaches in this vital field.

## 2 METHODS

### 2.1 Open Associative-memory, water-mediated, structure and energy model (OpenAWSEM)

In this study, openAWSEM is employed—a python implementation of the AWSEM protein coarse-grained force field devised by Wolynes and colleagues, as outlined by Lu et al.^41^ This innovative simulation framework is constructed on openMM^48^ to fulfill a rapid (GPU-enabled), adaptable, and user-friendly purpose. OpenAWSEM follows the legacy of the Associative-memory, Water-mediated, Structure and Energy Model (AWSEM) for molecular dynamics (MD) simulation, as introduced by Davtyan et al,^49^ enabling precise and efficient modeling of protein dynamics.

To facilitate understanding for all readers, within the AWSEM framework, each amino acid residue is represented by three atoms: C_α_, C_β_, and O, with glycine as the sole exception. The physicochemical characteristics of diverse side chains manifest in the C_β_ atoms. The AWSEM-MD simulation protocol has been applied across a spectrum of biological studies, from protein structure prediction,^49–52^ prognostication of protein binding events,^50^ investigations of protein aggregation phenomena,^46,47^ to the exploration of intricate protein-DNA assemblies and remodeling processes.^53–56^ For further details, the AWSEM-Suite web server offers additional resources for protein structure prediction.^57^

In the simulations described, openAWSEM utilizes a coarse-grained scheme where each integration step is precisely 5 femtoseconds (fs). This fine temporal resolution is critical for capturing rapid molecular interactions accurately. Each step advances the simulation by this small time increment, allowing for detailed control over the simulation’s temporal progression. Simulation trajectories were generated using the Langevin dynamics method.

To estimate the effective time covered by the simulation, the output frequency and the stride length are key factors. For example, if the output frequency is set to 1000 steps and the stride for analysis is similarly 1000 steps, this implies that data is recorded and analyzed at these intervals. In a scenario where the x-axis in **Figure 2B** spans 10^6^ steps, and considering the simulation scale, the effective time for this output is calculated by multiplying the number of steps (100) by the time per step, resulting in a total of 0.5 microseconds (µs). This calculation is based on 100×5×10^−3^ µs per step, effectively summing up to 0.5 µs for detailed analysis. This detailed temporal mapping is essential for a thorough understanding of the dynamics at play, providing critical insights for accurately modeling complex molecular behaviors over time.

### 2.2 Structural preparation and simulation protocol for Abeta monomer attaching fibrils

We used the CreateFibril2.5 tool^58^ to construct the fibril model of 62 chains. The program was used to replicate and extend selected protein chains from a specified PDB ID: 2MXU. This process involves setting various parameters to control the replication, rotation, and placement of monomers along the fibril axis. We used this tool to generate fibril models with various morphological features, e.g., straight, twisting. **Figure 1** presents the filamentous models generated using the tool. See Supporting Information in Figure S1 for additional examples.

**Figure 1.**
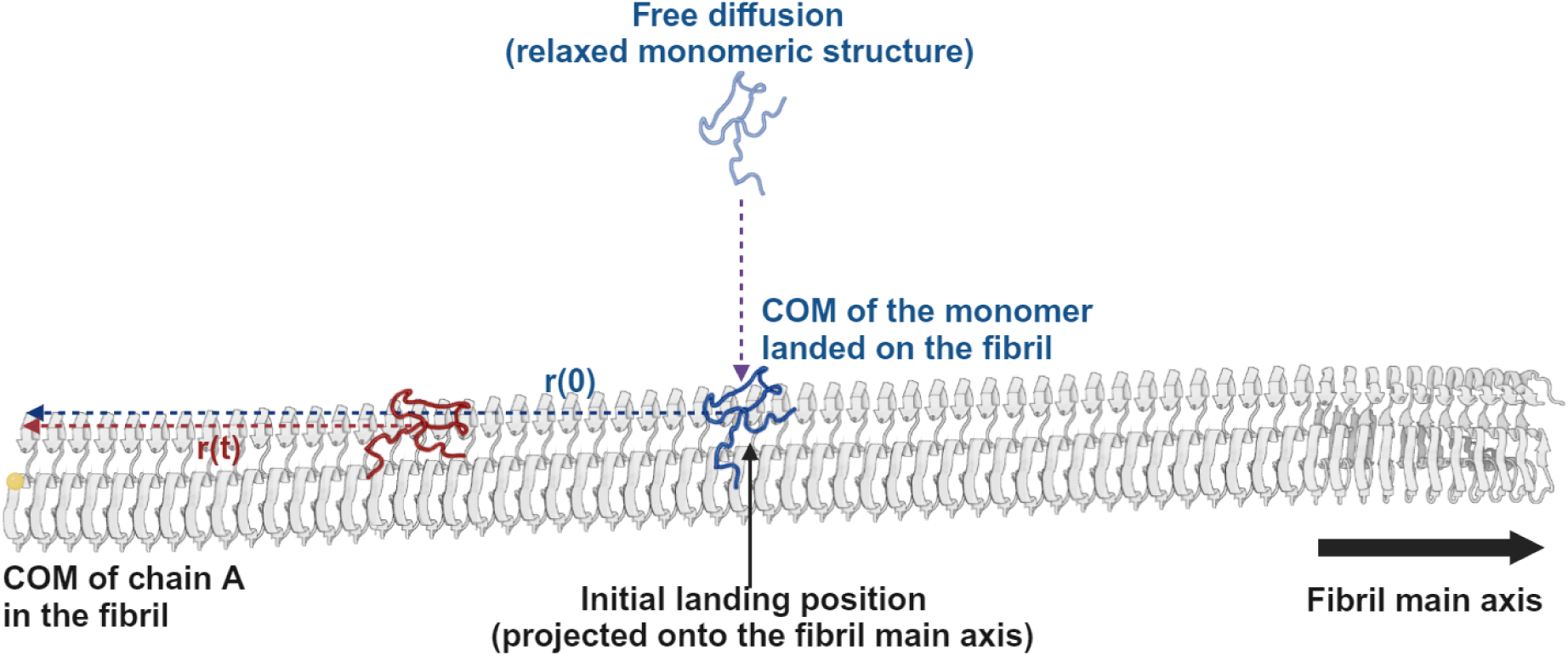
Schematic Diagram of Abeta Monomer Diffusion Simulation on Fibril Surfaces. This diagram illustrates a fibril filament consisting of 62 chains in a gray cartoon representation, alongside an Abeta monomer depicted in three scenarios: 1) Initially detached and freely diffusing with a relaxed structure, shown in transparent blue. 2) At the initial attachment point, *r*(0), highlighted in blue. 3) At its subsequent position, *r*(*t*), at time *t*, shown in red. The position vector, *r*, is defined from the center of mass (COM) of chain A, which is the leftmost chain in the fibril marked by a solid yellow circle, to the projected COM of the monomer along the fibril’s main axis (indicated by a thick arrow).

### 2.3 Nematic Order Parameter

To delineate the distinction in structure among fibril polymorphs, we utilized the nematic order parameter (P_2_ value) to quantify the structured arrangement on the surface of fibrils. Initially devised for delineating structural order in liquid crystals, the nematic order parameter was introduced in the investigation of protein aggregation by Caflisch and colleagues.^59^ Subsequently, this parameter was employed to elucidate aggregation of Amyloid peptides^42,60,61^ and has recently undergone comparison with order parameters acquired through neural network learning, as demonstrated by Charest et al.^62^ All these demonstrate its growing importance in the study of protein aggregation.

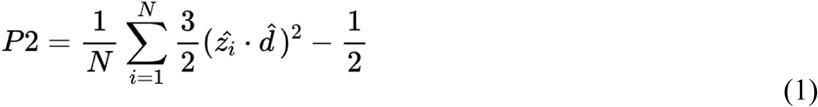

where *d̂* (the director) is a unit vector representing the preferred direction of the fibril surface (defined using residue 24 and 25 from each chain in the fibril), 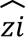 is the defined molecular vector (derived from residue 8 to 13 of the monomer), and *N* refers to the number of molecules considered in the calculation. Eq.(1) outputs a value between −0.5 and 1, where 1 indicates perfect alignment along the reference axis, 0 indicates a random orientation, and −0.5 indicates perfect perpendicular alignment. Values within this range reflect varying degrees of alignment. This method enables precise quantification of orientation differences by measuring the extent to which monomers align with or diverge from the fibril axis. It clearly distinguishes between various orientations and their impact on the diffusion process.

### 2.4 Mean Square Displacement

We use the mean square displacement (MSD) formula to calculate the diffusion coefficient (*D*) in a one-dimentional case, as shown below.

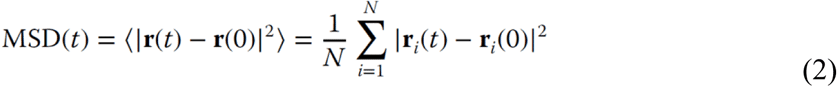

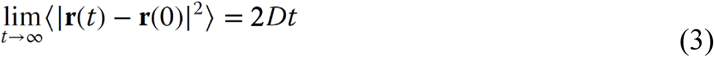

where *t* refers to time; *r*(0) represents the initial center-of-mass (COM) distance between the monomer (at *t*=0) and chain A (reference) in the fibril; *r*(*t*) indicates how the COM distance between the monomer and chain A in the fibril changes over time at *t*; *N* refers to the number of trajectories. To maintain accuracy and minimize deviation in vector calculations, the position vector *r* is defined as the distance from the center-of-mass (COM) of chain A (the leftmost chain in the fibril as the reference) to the projected COM of the monomer along the fibril’s main axis. By applying Eq.(2) to individual trajectories and then averaging the results, we can derive the diffusion coefficient as specified in Eq.(3), utilizing what is termed ‘trajectory-based’ approach for our analysis.

In scenarios where simulations capture monomer conformational or orientational changes, the diffusion dynamics can exhibit significant variability and may transition among different diffusion states. To accurately assess these dynamics, we segment each trajectory into smaller sections according to their orientations. These segments are then analyzed separately to calculate the diffusion coefficients specific to each orientation, employing what we refer to as a ‘segment-based’ approach. The modified diffusion coefficient is thus calculated as

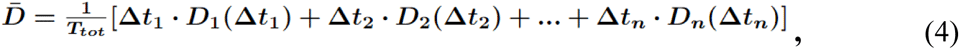

where 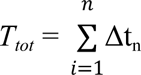 represents the sum of total time duration of trajectory segments; *D̅* is the overall diffusion coefficient; *D_n_* is the diffusion coefficient for n-th segment; Δt_n_ is the corresponding time duration. Eq.(4) provides a precise measure of how monomer’s conformation and orientation influence molecular mobility. Consequently, when calculating the diffusion coefficient, we account for the transitions between different monomer orientations within a single trajectory. By separating the trajectory based on each orientation and calculating the diffusion coefficient for each segment individually, we then apply a weighted average based on the frequency of each orientation. This approach ensures that the resulting average diffusion coefficient accurately represents the monomer’s true diffusion behavior.

### 2.5 Free energy profile for Abeta monomer diffusion on fibril surfaces

To understand the kinetics of the monomer diffusion, we computed the free energy of protein moving along fibril surfaces, specifically the potential of mean force (PMF), using the pyEMMA Python package, developed by Noe and collaborators.^63^ Calculating PMFs requires selecting a specific progress coordinate for sampling. Due to the difficulty of accessing high-energy configuration spaces through thermal activation alone, we employed a biasing force, a technique known as ‘importance sampling,’ to steer the sampling toward the desired configurational space. Our goal was to extensively sample the monomer distribution at various positions along the fibril’s main axis and to reweight these distributions to map the free energy landscape of protein diffusion accurately. This approach allows us to explore how spatial variations along the fibril affect energy barriers and stability.

For practical implementation, similar to previous studies, we conducted all sampling tasks using molecular dynamics simulations on the openAWSEM platform.^42^ To monitor the trajectory of a monomer as it diffuses towards a specific position on the fibril’s surface, we established numerous sampling windows. We applied a series of harmonic biasing restraints between the centers of mass of the monomer and the even end of the fibril. These restraints were set at intervals of approximately 2.2 Å, ranging from 20 to 230 Å from the even-end (chain A) of the fibril, positioning the monomer on the surface. This setup resulted in 110 independent simulations.

The biasing coordinate was defined as the center of mass distance between the monomer and chain A of the fibril, with a biasing force constant of 6 kcal/mol (∼25 kJ/mol) applied across all 110 simulations. Each simulation ran for 5 million time steps, outputting 5,000 frames for analysis at every 1,000 time steps. To manage data efficiently, we used a stride of 100, reducing the number of frames from 5000 to 50 for each simulation window. The collected data from these simulation trajectories, spanning 110 sampling windows, were then reweighted using the WHAM technique,^64^ as implemented in the pyEMMA package (thermo.wham).^63^ The distribution across all sampling windows is detailed in the Supporting Information Figure S2.

### 2.6 Surface Roughness (S_R_ value)

To assess the roughness of the fiber surface, we defined a reference plane that minimizes the vertical distance deviations of selected residue’s C_α_ atoms from it. First, we gather the 3D coordinates of all C_α_ atoms and calculate their centroid, which serves as the central reference point. The centroid represents the geometric center of the C_α_ atoms: residues 1 to 16 for N-terminal surface and residues 36 to 42 for C-terminal surface. Using this centroid, we compute the covariance matrix based on the positions of each atom relative to the centroid. This captures how the atoms are distributed in the *x*, *y*, and *z* directions. Specifically, we start with the coordinate matrix of full atoms, *X*,

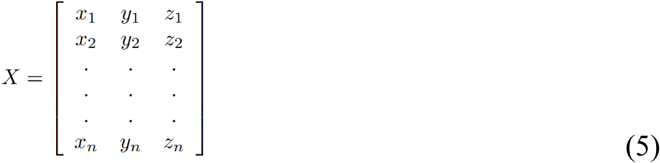

where n = 992, and the 3D coordinates of each atom is listed sequentially. The centroid of the atoms of the plane, **C**, is thus given by

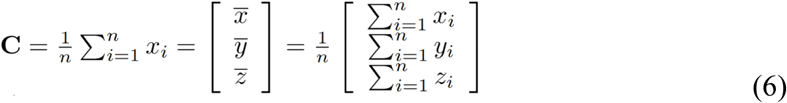

where x_i_ = [*x_i_*, *y_i_*, *z_i_*] are the coordinates of the *i*-th atom. At this step, we obtain the centroid as **C** = [*x̅*, *y̅*, *z̅*]. For the N-terminal surface, the calculated centroid is **C**_N_ =[−2.801, 0.040, 1.042], and for the C-terminal surface, the centroid is **C**_C_ =[10.205, −1.949, −3.695].Using the definition, we proceed to formulate the covariance matrix, accordingly

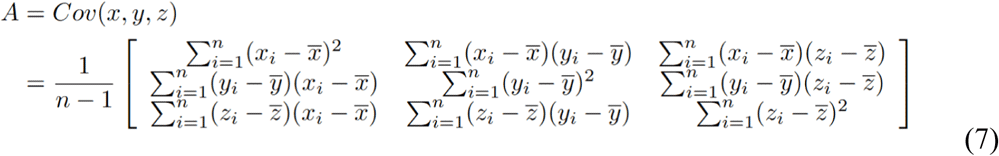

where *A* equal to *Cov(x, y, z)* represents the covariance matrix of the plane.

We then perform eigenvalue decomposition on the covariance matrix and obtain both eigenvalues and their corresponding eigenvectors,

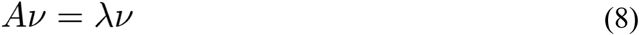

where *A* is the covariance matrix, *v* is eigenvector, and λ is the corresponding eigenvalue. The eigenvector associated with the smallest eigenvalue indicates the direction of least variation in the data—essentially the normal (perpendicular) direction to the plane we want to define (used to derive the coefficients of a plane equation, *a*x + *b*y + *c*z + *d* = 0). We obtain the normal vector as *n* = [*a*, *b*, *c*]. By positioning this normal vector to pass through the centroid, we obtain the optimized plane,

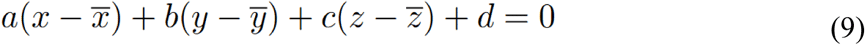

where *d* = -(*ax̅*+*by̅*+*cz̅*). Eq.(9) ensures that the calculated normal vector passes through the centroid. For the N-terminal surface, the values are *a*_N_ *=* −0.876, *b*_N_ *=* 0.229, *c*_N_ *=* 0.425, and *d*_N_ *=* −2.905. For the C-terminal surface, the values are *a*_C_ *=* 0.435, *b*_C_ *=* 0.806, *c*_C_ *=* 0.402, and *d*_C_ *=* −1.385. Figure 5A illustrates a schematic of this optimized plane intersecting the fibril surface structure. The root mean square (RMS) distances are calculated from the C_α_ atoms of residues 1 to 16 (N-terminal surface) or 36 to 42 (C-terminal surface) to this plane. It provides a quantitative measure of the structural deviation, which we define as surface roughness (S_R_ value).

## 3 RESULTS

### 3.1 Differential Diffusion Behaviors of Abeta Monomers Based on Orientation on Fibril Surfaces

In our study of Abeta monomers on fibril surfaces, we have identified significant variations in diffusion behaviors that are directly influenced by the monomers’ molecular orientation. To probe these variations more thoroughly, we conducted extensive simulations that illuminate the complex interplay between molecular structure and diffusion kinetics.

Expanding our understanding of these dynamics, **Figure 2** provides a crucial observation: it shows that a monomer on the C-terminal fibril surface can adopt two distinct orientations—parallel and perpendicular, as illustrated in **Figure 2A**. The Mean Squared Displacement (MSD) graph, plotted over time, furnishes vital insights into this diffusion phenomenon. By analyzing the slope of this graph, we determine the monomer’s diffusion coefficient in a one-dimensional (1-D) case. The diffusion coefficient analysis reveals that the orientation of the monomer significantly influences its diffusion dynamics on the surface. Specifically, our results indicate that monomers oriented parallel to the fibril main axis diffuse faster, exhibiting a larger diffusion coefficient (*D* (para.) = 19.9 μm²/s), compared to those oriented perpendicularly (*D* (perpend.) = 0.88 μm²/s) (illustrated in **Figure 2B**). To more comprehensively analyze the effect of monomer orientation on diffusion, we employed the P_2_ order parameter. This quantitative measure represents the degree of alignment of monomers relative to the fibril’s main axis (See METHODS for details). This method precisely quantifies orientation differences by assessing the alignment of monomers relative to the fibril main axis. It clearly distinguishes their effects on the diffusion process. **Figure 2C** presents P_2_ values across simulation time steps for three example trajectories. One trajectory with parallel orientation (in dark green) shows P_2_ values ranging from 0.6 to 1.0 (mean ∼ 0.8), while another in perpendicular orientation (in light green) ranges from −0.3 to −0.5 (mean ∼ −0.4). These trajectories exemplify the distinct orientations maintained throughout the simulation (See Supporting Information Figure S3, S4 for details). Notably, a significant orientational switching event is observed in a selected trajectory (in red), where P_2_ dramatically shifts from 0.8 to −0.3 at later simulation time steps (∼130 x 10^6^). Our simulations highlight several transitions from parallel to perpendicular orientations (See **Table 1**), indicating dynamic switching in monomer alignment that merits further exploration. **Figure 2D** offers a detailed analysis of the orientation distribution across all trajectories as the monomer diffuses along the fibril surface. Two predominant distributions emerge, corresponding to parallel and perpendicular orientations, centered around P_2_=0.8 and P_2_=−0.4, respectively. The average P_2_ values for these orientations, along with their associated error bars, are clearly depicted in the inset of **Figure 2D**.

**Figure 2.**
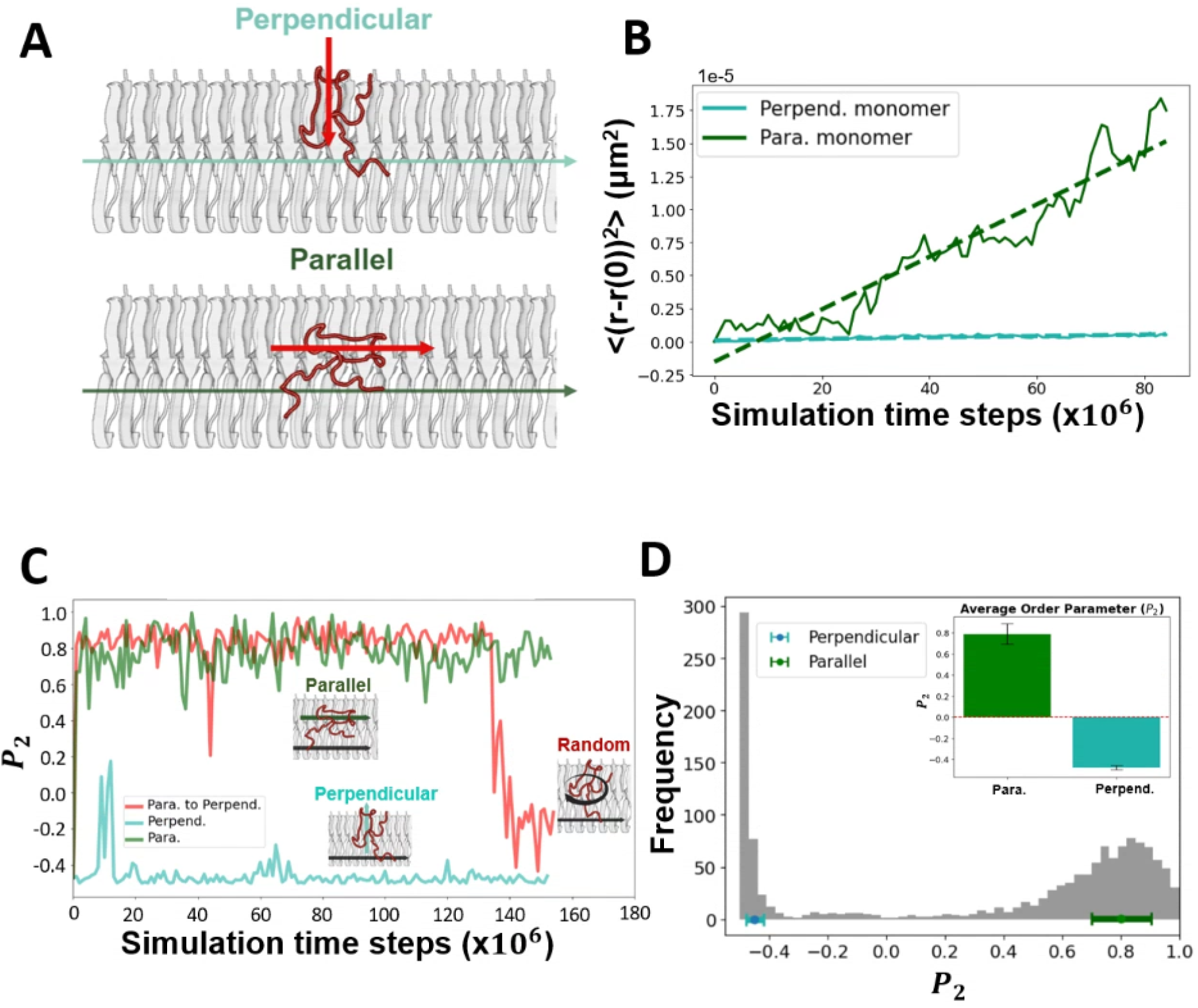
Analysis of an Abeta42 monomer’s orientation and its diffusion dynamics on the C-terminal fibril surface is presented. (A) A schematic diagram shows the monomer’s two main orientations—parallel and perpendicular—on the fibril surface. (B) Mean-squared displacement (MSD) over time of the entire trajectories (trajectory-based), with the diffusion coefficient calculated from the MSD using Eq. (2). The diffusion coefficients are calculated: *D* (para.) = 19.9 μm²/s and *D* (perpend.) = 0.88 μm²/s) (C) Changes in the order parameter (P_2_) for monomers with different orientations over simulation time steps. P_2_ values of ‘1’, ‘0’, and ‘-0.5’ correspond to ‘parallel’, ‘random’, and ‘perpendicular’ orientations, respectively, as illustrated in the inset. Three example trajectories are shown: 1. Parallel trajectory in dark green 2. Perpendicular in light green 3. Parallel-to-perpendicular trajectory in red. (D) Distribution of monomer orientation on the fibril in terms of P_2_. Inset: Comparison of the average order parameter P2 (Parallel vs Perpendicular); the perpendicular monomer is indicated in light green and the parallel monomer in dark green.

**Table 1.** Summary of Simulation Data Collection and Analysis Across Diverse Trajectories.

| Trajectory based <sup>a</sup> |  | 0 | 1 | 2 | 3 | 4 | 5 | 6 | 7 | 8 | 9 |
| --- | --- | --- | --- | --- | --- | --- | --- | --- | --- | --- | --- |
|  | Para. | ☑ |  |  | ☑ | ☑ | ☑ | ☑ | ☑ | ☑ |  |
|  | Perpend. |  | ☑ | ☑ |  |  |  |  |  |  | ☑ |
| # of orientation change |  | 1 | — | — | — | 1 | — | — | — | — | 1 |
| Segment based |  |  |  |  |  |  |  |  |  |  |  |
| Order parameter (P<sub>2</sub>) | Para. | 0-138 <sup>b</sup><br>(frame) | — | — | 0-153 | 0-90 | 0-100 | — | 0-100 | 0-100 | — |
|  | Perpend. | 139-154 | 0-152 | 0-154 | — | 91-152 | — | — | — | — | 0-50 |
| Diffusion coefficient | Para. | 85-137<br>(frame) | — | — | 15-153 | 55-106 | 15-100 | — | 55-100 | 15-100 | — |
|  | Perpend. | 140-154 | 15-152 | 15-154<br>(87-106) <sup>c</sup> | — | 107-152 | — | — | — | — | 35-60 |

<sup>a</sup>Checked box represents the primary orientation of the monomer during diffusion on the fibril surface.
<sup>b</sup>(*m-n*) represents the calculation is performed on frame '*m*' to frame '*n*'.
<sup>c</sup>(*m-n*) represents the second trajectory exhibited unusual P<sub>2</sub> values between frame *m* and frame *n*; therefore, its diffusion coefficient was calculated separately.

This analysis demonstrates that parallel-oriented monomers diffuse up to two orders of magnitude faster than their perpendicular counterparts, aligning with previously recorded experimental rates of 0.5 - 2.4 μm²/s, obtained via single-particle tracking with AFM on synthetic fibril surfaces.

Significantly, this research introduces the first computational model that computes peptide diffusion coefficients on fibril surfaces with results that closely match experimental observations. This breakthrough enhances our understanding of peptide behavior on fibrillar substrates and provides a robust framework for future experimental validation.

### 3.2 Monomer displays diverse diffusion states

Monomers can undergo orientation changes while diffusing on the fibril surface. To accurately quantify their 1-D diffusion coefficients, we meticulously analyze all simulation trajectories, categorizing them by the monomers’ orientations—either perpendicular or parallel. This process involves segmenting the trajectories into distinct sections based on orientation, resulting in numerous well-defined segments. We then calculate the diffusion coefficients for each segmented trajectory. In other words, the ‘segment-based’ analysis isolates the effects of orientation from those caused by orientation-switching events and conformational changes during monomer diffusion on the fibril. **Figure 3** illustrates our findings that even within the ‘parallel’ orientation, significant variations exist. We identified ‘fast’ and ‘slow’ diffusion states, with diffusion coefficients of 68.8 μm²/s and 7.66 μm²/s, respectively. This differentiation highlights the complexity of monomer dynamics on fibril surfaces (See Supporting Information Figure S5 to S7 for details). Table 2 provides a summary and comparison of the diffusion coefficients determined by both the trajectory-based and segment-based approaches.

**Figure 3.**
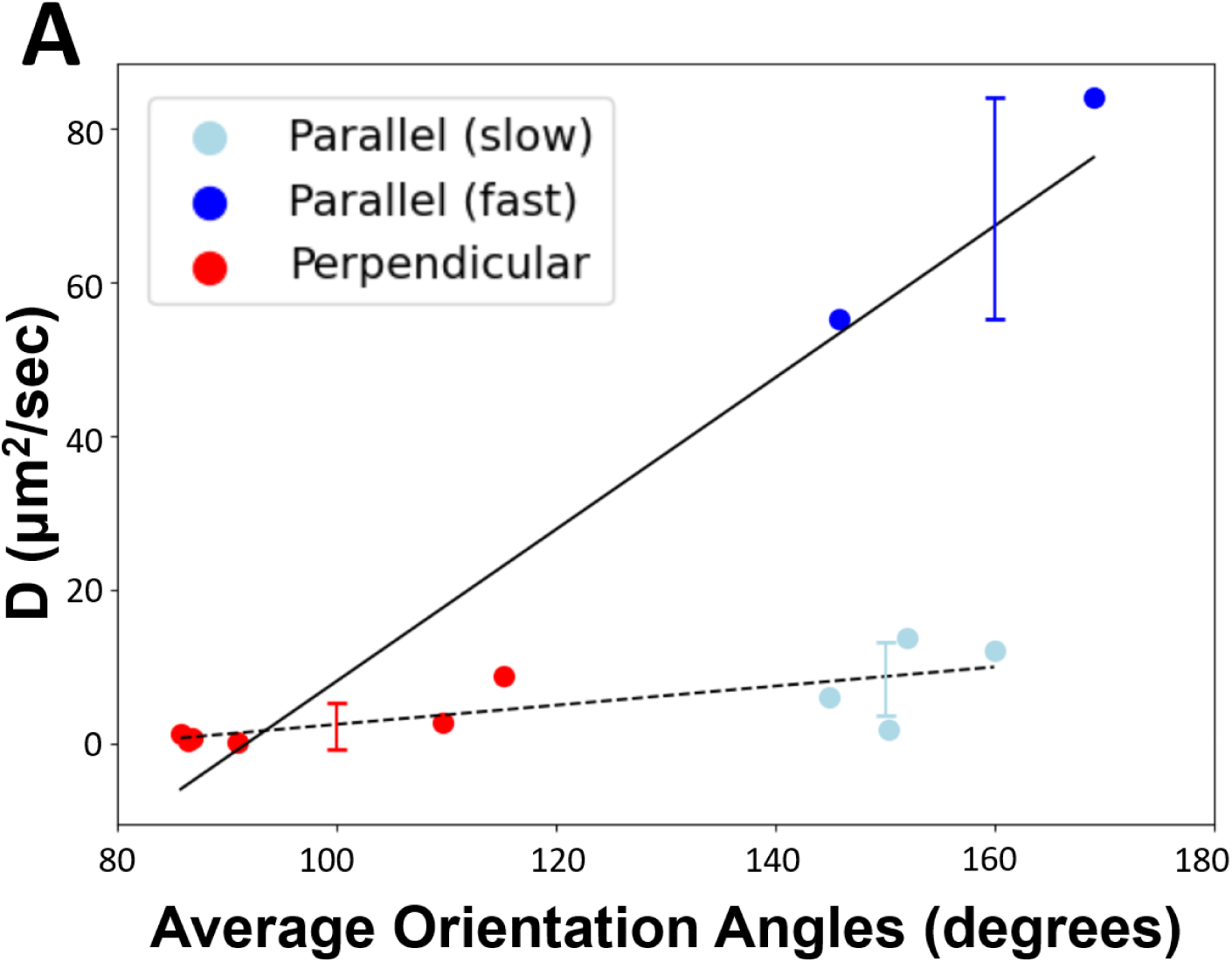
Further investigation of Abeta42 monomer’s multiple diffusion states on the C-terminal fibril surface is shown (using a segment-based approach) The diffusion coefficients calculated from eleven trajectory segments are represented by solid dots and are categorized into three distinct groups based on their diffusion dynamics and orientation, with blue dots indicating parallel orientation (two subgroups: light blue and regular blue) and red dots indicating perpendicular orientation. Monomers aligned parallel (140-170°) to the fibril, in general, diffuse faster (*D*_fast_^(*∥*)^ = 68.8 μm²/s; D_slow_^(*∥*)^ = 7.66 μm²/s) compared to those aligned perpendicularly (80-120°), which show slower diffusion rates (*D*^(⟂)^ = 0.88 μm²/s). The correlation between a monomer’s orientation and its diffusion coefficient on the fibril surface is clearly demonstrated, revealing two distinct diffusion states within the parallel-oriented monomers.

**Table 2.** Summary of Diffusion Coefficients Calculated.

| Trajectory-based approach |  | Para.<sup>(\|\|)</sup> | Perpend.<sup>(⊥)</sup> |  |
| --- | --- | --- | --- | --- |
| Diffusion coefficient (D) | C-ter | 19.9<br>($\mu\text{m}^2/\text{s}$) | 0.88 | |
|  | N-ter | 0.35 | — |  |
| Periodic length<br>(Free energy profile) | C-ter | 101<br>($\text{\AA}$) | 84 | |
| Segment-based approach |  | Para.<sub>fast</sub><sup>(\|\|)</sup> | Para.<sub>slow</sub><sup>(\|\|)</sup> | Perpend.<sup>(⊥)</sup> |
| Diffusion coefficient (D) | C-ter | 68.8<br>($\mu\text{m}^2/\text{s}$) | 7.66 | 0.88 |

### 3.3 Unveiling Asymmetry in Surface Diffusion along the Fibril: Computational Insight into Surface-mediated Fibril Growth

Our study explore the binding affinities of fibril surfaces and their position-dependent characteristics to better understand the mechanism of fibril growth kinetics. Using umbrella sampling, we explored the free energy landscape of an Abeta monomer on the C-terminal surface (See Motheds for details). We observed a notable decrease in free energy as the monomer approached a specific fibril end, indicating a preference for binding to the even end over the odd end. **Figure 4** illustrates the binding free energy of a monomer as a function of its position along the C-terminal fibril surface. The graph compares the ‘parallel’ and ‘perpendicular’ monomers with the odd end set as the energy reference point. We note that the overall free energies for both orientations (either parallel or perpendicular) declined towards the even end (left-hand side), punctuated by oscillatory fluctuations (will be discussed later). Notably, the even end of the simulated fibril system, approximately 25 nm away from the odd end, demonstrates a binding energy advantage of about 3-6 kcal/mol over the odd end. This observation aligns with the findings of a recent experimental study by Dev Thacker et al. (refer to Figure 5 in their publication)^65^, which reported asymmetric growth at the fiber endpoints. Our computational results support the experimental evidence of fibril asymmetry and offer deeper insight by showing that the even end has a lower free energy.

**Figure 4.**
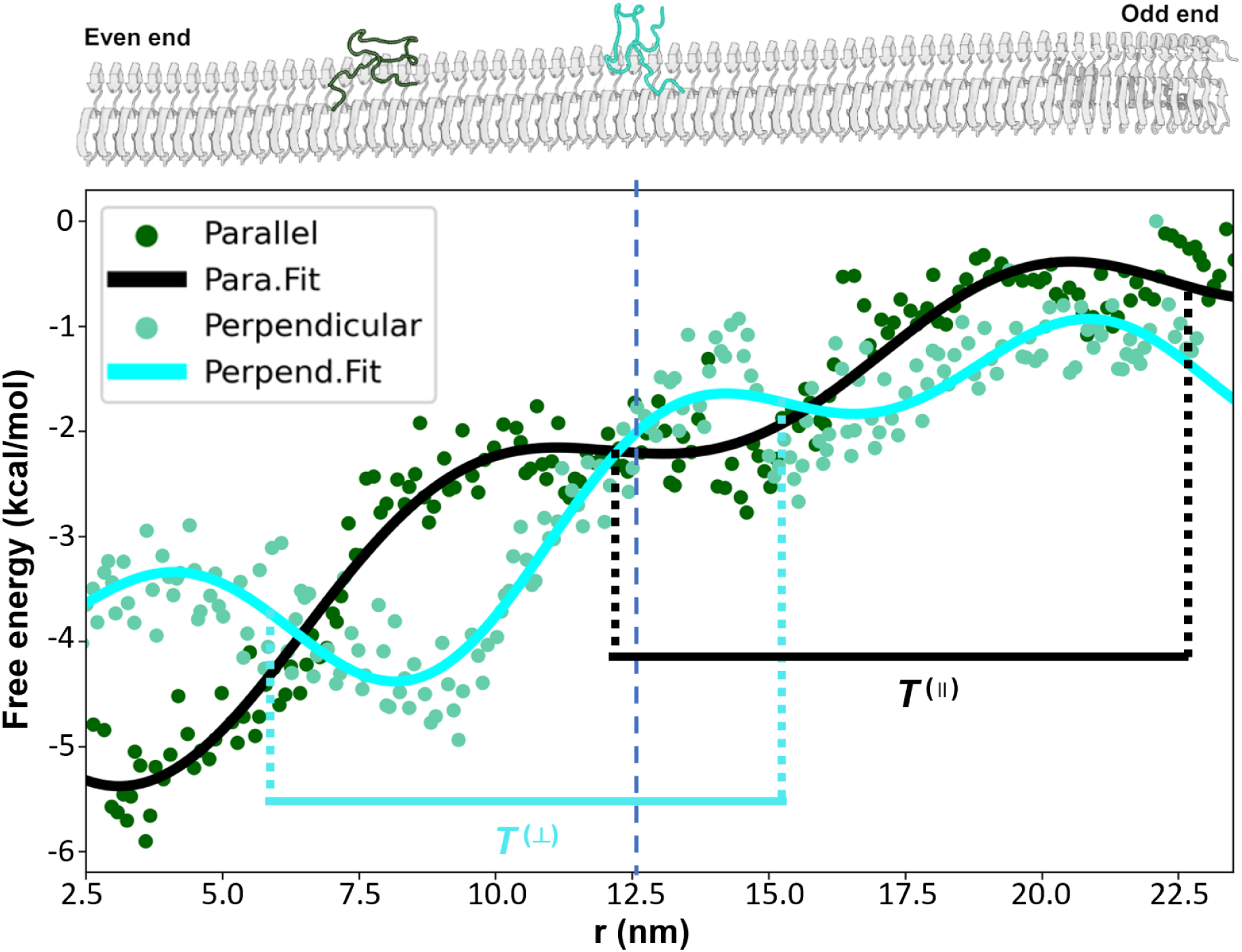
The diffusion free energy profiles of an Abeta42 monomer with ‘parallel’ and ‘perpendicular’ orientations on the C-terminal fibril surface are shown. This profile is plotted as a function of the monomer’s position along the fibril surface, extending from the even end to the odd end. Two orientations of the monomer are examined: dark green dots represent the free energy of monomers oriented parallel to the surface. In contrast, light green dots indicate the free energy of monomers oriented perpendicularly. The solid curves are the corresponding fit curves with their periodic length indicated by *T*^(∥)^ (parallel) and *T*^(⟂)^ (perpendicular);. The accompanying diagram of the fibril structure, shown above, clearly marks the even end on the left and the odd end on the right.

Additionally, our simulation uncovers a previously unreported oscillatory behavior in free energy, as evidenced by the fitted curves of the data. This oscillation enables monomers aligned parallel to the fibril axis to exhibit additional stabilization or destabilization energy, ranging ± 2 kcal/mol, compared to those in perpendicular orientations. The magnitude of this energy variation is contingent upon the monomer’s position along the fibril. For instance, at the 4 nm position, a monomer in a parallel orientation displays a free energy that is approximately 2 kcal/mol more stable than a monomer in a perpendicular orientation. Conversely, at the 9 nm position—just 5 nm away—the same parallel monomer becomes about 2 kcal/mol less stable compared to its perpendicular counterpart. Furthermore, we analyzed the oscillatory behavior of free energy and computed the periodic length for both parallel and perpendicular orientations. Our findings reveal that the parallel orientation exhibits a periodicity of approximately 101 Å, roughly equivalent to the length of 24 Abeta chains in the fibril, with each chain spanning an interval of about 4.2 Å. In contrast, the perpendicular orientation shows a periodic length of about 84 Å, which corresponds to the length of 20 Abeta chains. This discovery is pivotal as it illuminates the complex mechanics of surface-mediated fibril growth, providing a more comprehensive understanding of the processes that govern this phenomenon.

### 3.4 Quantitative Surface Roughness Analysis Clarifies Differences Between N-terminal and C-terminal Surfaces

Up to this point, our results have demonstrated that the diffusion dynamics of the Abeta monomer depend on its orientation, particularly on the C-terminal (C-ter) fibril surface. To expand our understanding and explore the effects of different fibril surface types, we conducted a comparative analysis, focusing particularly on the N-terminal (N-ter) surfaces, as depicted in **Figure 5**. The N-ter surface of the fibril structure exhibits an inward curved, groove-like shape. The groove provides an environment that enhances the formation of physical contacts with the monomer. As shown in **Figure 5A**, this unique geometry causes the monomer to reposition its orientation to align parallel within the groove, affecting its diffusion dynamics. Consequently, as illustrated in **Figure 5B**, the diffusion coefficient (D) of the monomer on the N-ter surface tends to be lower compared to that on the C-ter surface, indicating a slower diffusion rate. (See Supporting Information Figure S8 for details)

**Figure 5.**
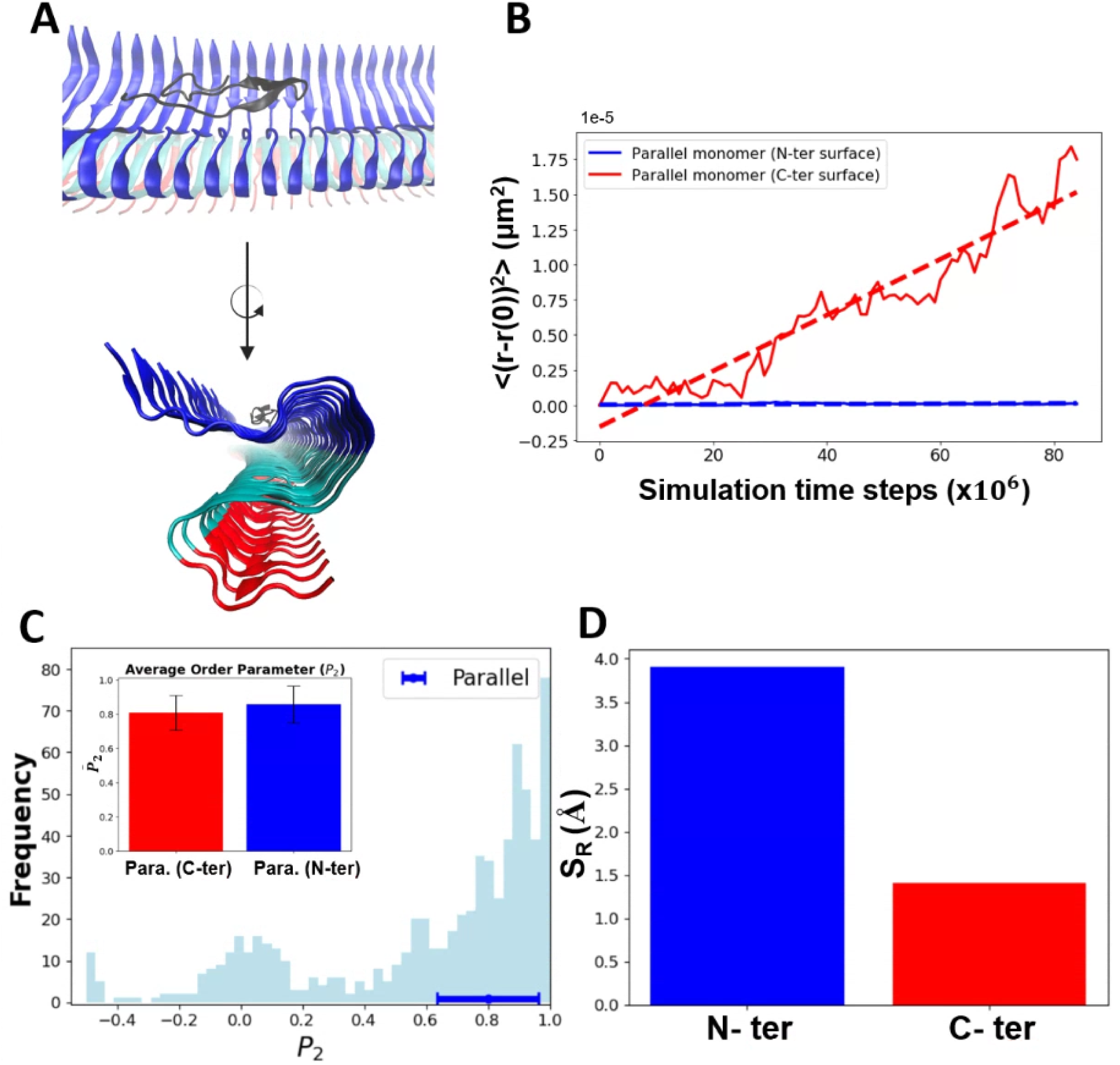
The variations in diffusion behaviors across different types of fibril surfaces (N-terminal vs C-terminal) are shown. (A) Structural diagram of a protofibril filament, where the N-terminal (N-ter) surface is depicted in blue, and the C-terminal (C-ter) surface is shown in red. A monomer in a ‘parallel’ orientation is depicted on the N-ter surface in black. Panel (B) compares the Mean Squared Displacement (MSD) analysis of the monomer in ‘parallel’ orientation on the C-ter surface, represented by the red line (diffusion coefficient *D* = 19.9 μm^2^/s), to the MSD on the N-ter surface, represented by the blue line (diffusion coefficient *D* = 0.35 μm^2^/s). Panel (C) features the distribution of the P_2_ value across the entire set of simulation trajectories. It highlights a major population exhibiting ‘parallel’ orientation at approximately P_2_=0.8, with its error bar indicated using blur color. The inset in this panel provides a comparison of the average P_2_ values of the monomer on the N-ter and C-ter surfaces, shown with blue and red bars respectively, including error bars. Panel (D) displays a bar chart that compares the surface roughness (S_R_) of the N-ter and C-ter surfaces. It illustrates that the surface roughness of the N-ter (shown in blue) is greater than that of the C-ter (shown in red).

**Figure 5C** presents the distribution of the P_2_ values across all simulation trajectories on the N-ter surface, where a significant population aligns around P_2_=0.8, indicative of a ‘parallel’ orientation (See Supporting Information Figure S4 for details). This result aligns with observations on both the N-ter and C-ter surfaces, confirming consistent orientation across different fibril regions, as illustrated in the inset in the figure. Interestingly, despite the monomers displaying similar parallel orientations, notable differences in their diffusion properties are observed. We hypothesize that these variations are due to the distinct geometric shapes of the N-ter and C-ter surfaces, which significantly influence their diffusion behaviors.

Our analysis extended beyond the P2 properties of the monomers to a deeper examination of the structural differences between the N-ter and C-ter surfaces. Structurally, the C-ter surface is smoother, offering a more favorable landing area for monomers, whereas the N-ter surface is comparatively rougher. To quantify this, we employ the average Root Mean Square (RMS) Distance, measuring the fluctuations in distance between C_α_ atoms and the best-fit plane. This optimized plane is defined by a normal vector corresponding to the smallest eigenvalue (termed as the Surface Roughness (S_R_) value). Conceptually, a higher RMS Distance indicates greater surface roughness due to increased fluctuations. Our RMS distance analysis confirms that the N-ter surface is indeed rougher than the C-ter surface, as illustrated in **Figure 5D. Figure 6A** illustrates the generation of an optimized plane and how to calculate the surface roughness accordingly (See Methods for details). **Figure 6B** compares the S_R_ value of N-terminal and C-terminal surfaces and shows the N-terminal surface’s higher S_R_ value. In conclusion, our findings suggest that the surface shape not only influences the monomer’s orientation but also entraps it in a state of reduced diffusive dynamics.

**Figure 6.**
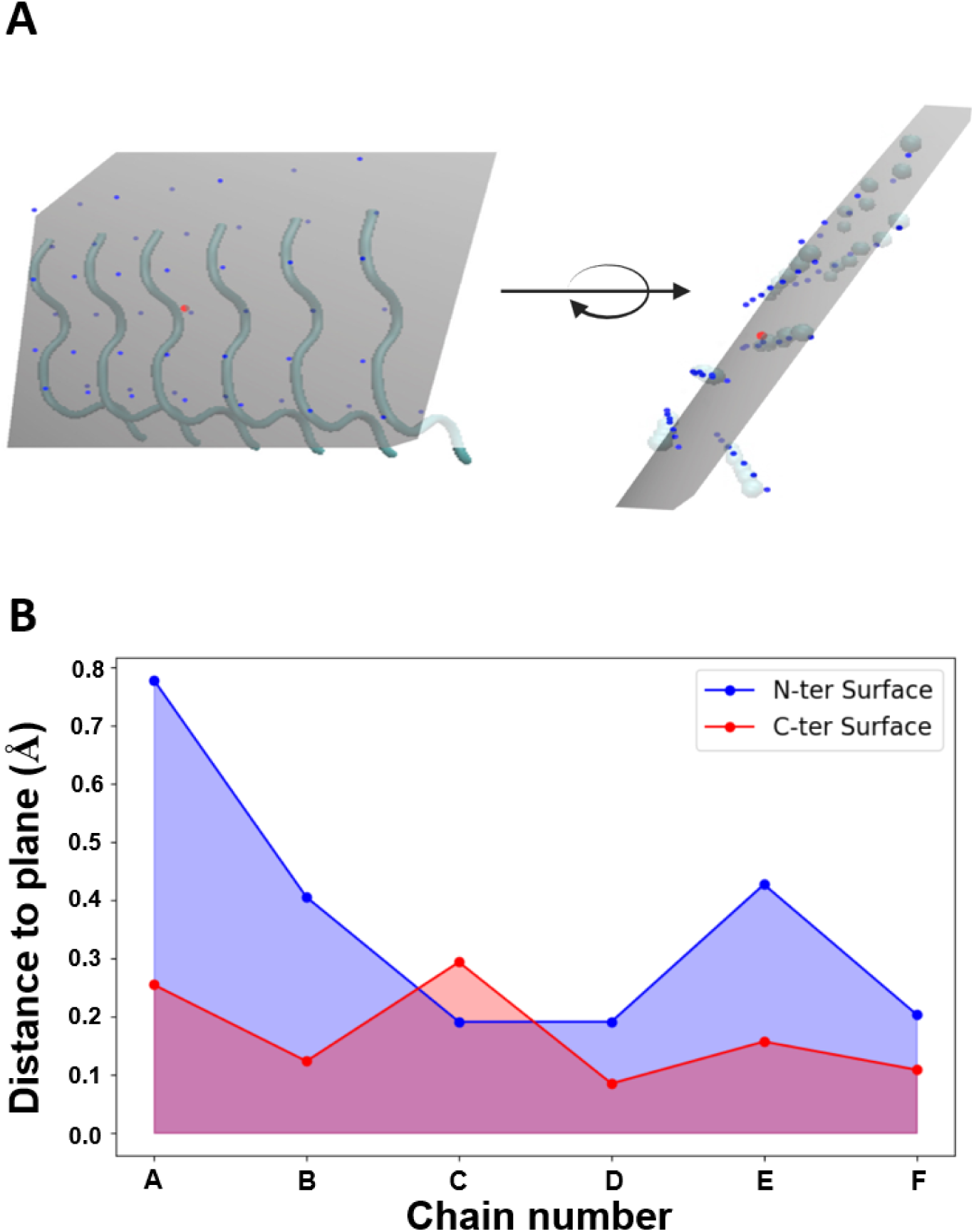
Schematic of the surface roughness calculation is presented. (A) An optimized plane (in gray) intersecting the fibril structure is used to calculate the RMS distances of the C_α_ atoms (solid blue circles) to the plane. The cartoon diagram represents the backbone structure of the peptides in the fibril; the C-terminal surface is used as the example. The solid red circle refers to the geometric center of the C_α_ atoms where the optimized plan intersects. A rotation angle is used to guide the visualization from different angle perspectives. (B) The surface roughness is shown as a function of the chain number in the fibril (chain A to F).

## 4 DISCUSSIONS

### 4.1 Monomer Orientation’s Active Role in Fibril Growth

It is suggested that the fibril surface contributes actively to fibril growth, rather than serving merely as a passive scaffold.^65,66^ This perspective offers a nuanced understanding of amyloid protein dynamics on pre-existing fibrils, an area critical for advancing our approach to neurodegenerative diseases.^23^ Investigations employing molecular dynamics (MD) simulations^60,67^ and High-Speed Atomic Force Microscopy (HS-AFM)^68^ have revealed how alterations in protein dynamics on fibrils correlate with protein misfolding, aggregation, and disease progression. These methods allow for detailed observation and analysis of the structural and dynamic changes in amyloid proteins, providing insights into their role in disease mechanisms. Deviations in these dynamic processes could lead to abnormal fibril growth and morphology, potentially culminating in cellular toxicity.^20^ A recent integrated theoretical study highlighted the role of diffusion dynamics of Abeta monomers on fibril surfaces, introducing ‘Bulk-docking’ and ‘Surface-docking’ as key pathways.^69^ This analysis reinforces the significance of their mechanisms in modulating the growth kinetics of amyloid-like fibrils.

In this study, we have identified three distinct states of monomer diffusion, influenced by their orientation, conformational changes, and interactions at the surface interface. These findings highlight how the orientation of the monomer affects its diffusion dynamics on the fibril surface. Specifically, monomers in a perpendicular orientation demonstrate slower diffusion rates (D_slow_^(⟂)^), while monomers in a parallel orientation exhibit both slower (D_slow_^(*∥*)^) and faster (D_fast_^(*∥*)^) diffusion states. The subscript in our notation specifies the diffusion rate as ‘slow’ or ‘fast’, and the superscript indicates the orientation—’perpendicular’ (⟂) or ‘parallel’ (*∥*). Furthermore, we have observed transitions between these diffusion states. This finding underscores the importance of orientation and dynamic states in controlling diffusion mechanisms.

Various polymorphic fibril types may exhibit distinct diffusion dynamics, influenced by surface heterogeneity.^70–72^ For instance, ‘helical’ fibrils could regulate diffusion rates by altering the orientation of monomers on their surface, such as by modulating sliding speeds, as observed from this work. These micro-level changes might have significant implications at the macroscopic level, possibly affecting the overall rate of fibril growth. A scenario where monomers slide excessively rapidly might be less favorable. That being said, nature avoids straight filamentous fibrils for biological use. To date, there are no experimental studies specifically showing how changes in monomer orientation on fibrils can alter the overall growth rate of the fibrils. However, our results suggest that fibrillar twisting can facilitate slower protein diffusion along the fibril surfaces (See Supporting Information in **Figure S9**), which favors elongation at rates compatible with biological timescales, as illustrated in **Figure 7**. Such a hypothesis is compatible with the view of fibril surface-mediated growth kinetics, as evidenced by both theoretical and computational studies.^42,69^ Our finding suggests that the physical structure and surface properties of fibrils are not just passive elements but actively influence the behavior and movement of amyloid proteins in fibril growth. Recent computational analyses reveal a complex ensemble structure at fibril tips, characterized by the collective motions of several peptide chains. These motions, rate-limiting for fibril elongation, involve overcoming significant non-native contacts.^73^ This indicates a critical interaction between the molecular-scale dynamics and the larger-scale structural behaviors, linking the diffusion of individual peptides on fibril surfaces to the emergent properties at fibril tips. Specifically, the collective behavior of peptides at the fibril tips, involving complex and coordinated movements, could be key to understanding how the physical and chemical characteristics of the fibril itself control fibril elongation. Such insights could be pivotal in understanding the mechanistic pathways involved in the progression of neurodegenerative diseases.

**Figure 7.**
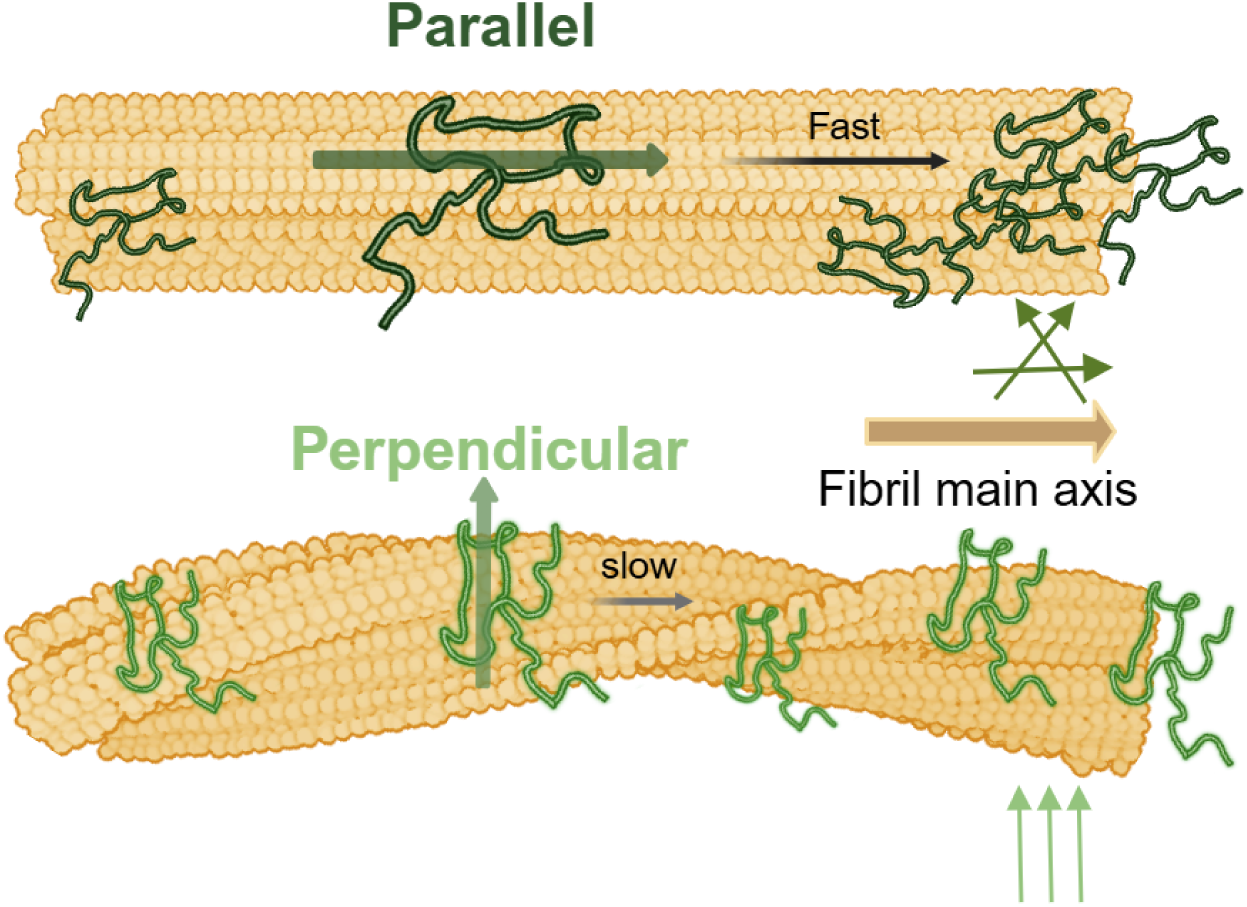
Schematic illustrating the influence of straight and twisted fibril structures on the preferred orientation of monomers. This figure illustrates the pivotal role of fibrillar twisting in controlling monomer diffusion dynamics through orientation modulation. The top schematic depicts rapid monomer diffusion toward one fibril end, leading to a disordered assembly of many monomers. This configuration may impact fibril growth. In contrast, the bottom schematic shows slower monomer diffusion, where accumulated monomers form an ordered assembly that facilitates efficient fibril growth within biological conditions. It’s important to note that the fibril formation discussed here does not necessarily relate to neurotoxicity, but rather to the type of fibril assembly that does not generate toxic oligomers.

### 4.2 Hypothesis on the Kinetic Stages of Fibril Growth via Surface-Mediated Processes

While traditional views on fibril elongation primarily focus on monomers bulk-docking at fibril tips, recent insights highlight the significant role of surface-mediated processes. Unlike the restricted, rod-like areas at the fibril tips, fibril surfaces provide extensive interaction zones, making surface-docking more advantageous than bulk-docking. This understanding is framed by a hypothesis that fibril growth involves a sequential three-stage process: 3D bulk-diffusional attachment of monomers to the fibril surface, 1D surface-mediated diffusion along the surface, and finally, a conformational conversion into the fibril structure.

#### Stage One: 3D Bulk-Diffusional Attachment

- Initially, monomers move from the bulk solution to the fibril surface. At low protein concentrations, the infrequent arrival of monomers on the surface may slow down the growth, potentially becoming the rate-determining step.

#### Stage Two: 1D Surface-Mediated Diffusion

- After attachment, monomers traverse along the fibril surface seeking optimal sites for integration into the fibril structure. The efficiency of this process becomes increasingly critical as protein concentrations rise, influencing the rate of fibril elongation.

#### Stage Three: Conformational Conversion into the Fibril Structure

- The final stage involves the monomers undergoing conformational changes to integrate into the fibril. The rate of this stage is influenced by local conditions and could become more significant at higher protein concentrations.

This comprehensive model suggests that at lower concentrations, bulk-diffusional attachment dominates the rate-determining process. However, as concentrations increase, the critical step may shift to surface-mediated diffusion or conformational conversion, depending on monomer availability and specific surface conditions. The extensive surface area of fibrils facilitates the attachment of multiple monomers, supporting a concentration-dependent elongation rate as observed at low Abeta concentrations (1-3 µM). This framework provides a mechanistic basis for the observed ‘stop-go’ elongation behavior, underscoring the complex interplay between kinetics and protein concentration in fibril growth.^74^

### 4.3 Asymmetric Fibril Ends: Influencing Monomer’s 1D Diffusion and Growth Dynamics

In studying amyloid fibrils, it is crucial to understand the asymmetric effects at the fibril ends due to their substantial influence on fibril growth and stability. Each end of the fibril displays unique kinetic and structural characteristics that dictate monomer addition and affect the rate of fibril elongation. Insights into these variations are critical for the development of targeted therapies that could selectively interact with the more reactive fibril end to potentially halt fibril extension.

Fibril growth demonstrates distinctive behaviors at the even and odd ends, driven by structural differences. At the even end, the Aβ peptide tends to adopt a closed conformation where the β1 and β2 sheets are closely juxtaposed, leading to minimal fluctuations. This stability is linked to the robust β-sheet interactions facilitated by the β1 sheet exposed at the even end.^75^ In contrast, at the odd end, the Aβ peptide is more variable, often adopting an open conformation with more widely spaced β-sheets. This is attributed to weaker β-sheet interactions due to the exposure of the β2 sheet at this end.^75^ These structural differences explain why fibril extension predominantly occurs from the odd end, which exhibits more dynamic structural states.^75,76^

While previous studies primarily observed fibril elongation at the odd end, indicating a kinetic preference for growth in this region, our research introduces a contrasting perspective by revealing a distinct sliding preference towards the even end based on energetic analyses. This suggests a thermodynamic favorability at the even end, offering a new viewpoint on fibril elongation dynamics that contrasts with earlier kinetic-focused studies. This divergence invites further investigation into the conditions under which thermodynamic versus kinetic factors dominate fibril growth, enhancing our understanding of amyloid fibril mechanics and providing novel targets for therapeutic intervention.

Our findings show that the binding free energy of a monomer varies along the fibril surface depending on its orientation—parallel or perpendicular—with the odd end serving as the reference. The free energy generally decreases towards the even end, accompanied by oscillatory profiles. Notably, the even end of our ∼25 nm fibril system demonstrates a binding energy advantage of 3-6 kcal/mol over the odd end. Monomers aligned parallel to the fibril axis additionally benefited from a stabilization/destabilization energy of about ± 2 kcal/mol compared to those in a perpendicular orientation. These results, when scaled per chain in the fibril, translate to 0.05-0.1 kcal/mol per amino acid for even over odd end, and ± 0.03 kcal/mol per amino acid for parallel over perpendicular alignments.

The oscillatory behaviors in free energy are believed to be linked to the surface properties at the monomer-fibril interfaces, including different types of residue-residue interactions, their resultant friction effects, and the size of the monomer itself. We have measured the monomer’s size in both parallel and perpendicular orientations along the fibril axis, finding sizes of 35.5 Å and 24.0 Å, respectively. Comparing these values with their corresponding free energy periods of 101 Å and 84 Å confirms the oscillatory nature of the free energy profile (or potential of mean force). This suggests that the diffusion dynamics are influenced by biased periodic potentials, akin to those experienced by a Brownian particle during its random walk process. To further underscore this analogy, we explored the verification of barrier-crossing events within these biased effective potentials, illustrating how they mimic the random walk dynamics and facilitate understanding of monomer diffusion on fibril surfaces. There may be a scaling relation suitable for these effective periodic potentials, a topic warranting further investigation. All these findings highlight the role of diffusion energy preferences due to fibril end asymmetry and support the theory of surface-mediated processes in fibril growth kinetics, bridging simulation results with theoretical models.

## 5 CONCLUSIONS

This study provides critical insights into the role of surface-mediated secondary processes in Alzheimer’s disease, particularly in the context of Abeta42 peptide aggregation. Through the use of coarse-grained molecular dynamics simulations, we focused on elucidating the molecular intricacies of these secondary processes, distinct from primary nucleation.

Central to our investigation was the analysis of a freely diffusing Abeta42 monomer in proximity to pre-formed fibril structures. We meticulously calculated the diffusion coefficient (*D*) of the monomer as it interacted with both straight and twisted fibril surfaces, taking into account different monomer orientations. Our research reveals a strong and consistent correlation between the monomer’s diffusion coefficient and its orientation on the fibril surface. This finding is crucial, as it remains valid across different degrees of fibrillar twisting, emphasizing the significance of monomer orientation in surface-mediated secondary aggregation processes. This is the first simulation work to reveal roles of monomer conformational dynamics in determining kinetic behaviors of its diffusion on fibril surfaces.

Additionally, our study explored the differential effects of fibril surfaces on monomer dynamics. We conducted a comparative analysis of the C-terminal and N-terminal surfaces, observing that monomers exhibit lower diffusion coefficients on the N-terminal surface. This variation in diffusion dynamics is largely attributable to differences in surface roughness, quantified using the surface roughness (S_R_). Such disparities in shape and roughness of the fibril surfaces are pivotal in shaping the monomer dynamics, thereby influencing the secondary aggregation process.

In conclusion, our findings highlight the complex and nuanced role of surface characteristics in the secondary aggregation processes of Abeta42 peptides in Alzheimer’s disease. By shedding light on the molecular dynamics of these processes, our study opens new avenues for understanding and potentially intervening in the progression of Alzheimer’s disease, focusing on the manipulation of surface-mediated secondary aggregation mechanisms.

## 8 ACKNOWLEDGMENT

This work was supported by the National Science and Technology Council (NSTC), Taiwan, Grant No. NSTC 112-2113-M-194-012 and NSTC 113-2628-M-194-001-MY3. We thank to National Center for High-performance Computing (NCHC) of National Applied Research Laboratories (NARLabs) in Taiwan for providing computational and storage resources.

## REFERENCES

(1) Hardy, J.; Selkoe, D. J. The Amyloid Hypothesis of Alzheimer’s Disease: Progress and Problems on the Road to Therapeutics. Science 2002, 297 (5580), 353–356.

(2) Knopman, D. S.; Jones, D. T.; Greicius, M. D. Failure to Demonstrate Efficacy of Aducanumab: An Analysis of the EMERGE and ENGAGE Trials as Reported by Biogen, December 2019. Alzheimers. Dement. 2021, 17 (4), 696–701.

(3) Center for Drug Evaluation; Research. FDA’s decision to approve new treatment for Alzheimer’s disease. U.S. Food and Drug Administration. https://www.fda.gov/drugs/news-events-human-drugs/fdas-decision-approve-new-treatment-alzheimers-disease (accessed 2023-12-23).

(4) Piller, C. Second death linked to potential antibody treatment for Alzheimer’s disease. Science News. https://www.science.org/content/article/second-death-linked-potential-antibody-treatment-alzheimer-s-disease?utm_source=sfmc&utm_medium=email&utm_campaign=DailyLatestNews&utm_content=alert&et_rid=504035807&et_cid=4509061.

(5) Bucciantini, M.; Giannoni, E.; Chiti, F.; Baroni, F.; Formigli, L.; Zurdo, J.; Taddei, N.; Ramponi, G.; Dobson, C. M.; Stefani, M. Inherent Toxicity of Aggregates Implies a Common Mechanism for Protein Misfolding Diseases. Nature 2002, 416 (6880), 507–511.

(6) Walsh, D. M.; Klyubin, I.; Fadeeva, J. V.; Cullen, W. K.; Anwyl, R.; Wolfe, M. S.; Rowan, M. J.; Selkoe, D. J. Naturally Secreted Oligomers of Amyloid Beta Protein Potently Inhibit Hippocampal Long-Term Potentiation in Vivo. Nature 2002, 416 (6880), 535–539.

(7) Kayed, R.; Head, E.; Thompson, J. L.; McIntire, T. M.; Milton, S. C.; Cotman, C. W.; Glabe, C. G. Common Structure of Soluble Amyloid Oligomers Implies Common Mechanism of Pathogenesis. Science 2003, 300 (5618), 486–489.

(8) Nguyen, P. H.; Ramamoorthy, A.; Sahoo, B. R.; Zheng, J.; Faller, P.; Straub, J. E.; Dominguez, L.; Shea, J.-E.; Dokholyan, N. V.; De Simone, A.; Ma, B.; Nussinov, R.; Najafi, S.; Ngo, S. T.; Loquet, A.; Chiricotto, M.; Ganguly, P.; McCarty, J.; Li, M. S.; Hall, C.; Wang, Y.; Miller, Y.; Melchionna, S.; Habenstein, B.; Timr, S.; Chen, J.; Hnath, B.; Strodel, B.; Kayed, R.; Lesné, S.; Wei, G.; Sterpone, F.; Doig, A. J.; Derreumaux, P. Amyloid Oligomers: A Joint Experimental/Computational Perspective on Alzheimer’s Disease, Parkinson’s Disease, Type II Diabetes, and Amyotrophic Lateral Sclerosis. Chem. Rev. 2021, 121 (4), 2545–2647.

(9) Oosawa, F.; Asakura, S. Thermodynamics of the Polymerization of Protein; Molecular Biology S.; Academic Press: San Diego, CA, 1975.

(10) Oosawa, F.; Kasai, M. A Theory of Linear and Helical Aggregations of Macromolecules. J. Mol. Biol. 1962, 4, 10–21.

(11) Lee, J.; Culyba, E. K.; Powers, E. T.; Kelly, J. W. Amyloid-β Forms Fibrils by Nucleated Conformational Conversion of Oligomers. Nat. Chem. Biol. 2011, 7 (9), 602–609.

(12) Xu, Y.; Safari, M. S.; Ma, W.; Schafer, N. P.; Wolynes, P. G.; Vekilov, P. G. Steady, Symmetric, and Reversible Growth and Dissolution of Individual Amyloid-β Fibrils. ACS Chem. Neurosci. 2019, 10 (6), 2967–2976.

(13) Yuan, M.; Tang, X.; Han, W. Anatomy and Formation Mechanisms of Early Amyloid-β Oligomers with Lateral Branching: Graph Network Analysis on Large-Scale Simulations. Chem. Sci. 2022, 13 (9), 2649–2660.

(14) Knowles, T. P. J.; Waudby, C. A.; Devlin, G. L.; Cohen, S. I. A.; Aguzzi, A.; Vendruscolo, M.; Terentjev, E. M.; Welland, M. E.; Dobson, C. M. An Analytical Solution to the Kinetics of Breakable Filament Assembly. Science. 2009, pp 1533–1537. 10.1126/science.1178250.

(15) Cohen, S. I. A.; Linse, S.; Luheshi, L. M.; Hellstrand, E.; White, D. A.; Rajah, L.; Otzen, D. E.; Vendruscolo, M.; Dobson, C. M.; Knowles, T. P. J. Proliferation of Amyloid-β42 Aggregates Occurs through a Secondary Nucleation Mechanism. Proc. Natl. Acad. Sci. U. S. A. 2013, 110 (24), 9758–9763.

(16) Bishop, M. F.; Ferrone, F. A. Kinetics of Nucleation-Controlled Polymerization. A Perturbation Treatment for Use with a Secondary Pathway. Biophys. J. 1984, 46 (5), 631–644.

(17) Ferrone, F. A.; Hofrichter, J.; Eaton, W. A. Kinetics of Sickle Hemoglobin Polymerization. II. A Double Nucleation Mechanism. J. Mol. Biol. 1985, 183 (4), 611–631.

(18) Ruschak, A. M.; Miranker, A. D. Fiber-Dependent Amyloid Formation as Catalysis of an Existing Reaction Pathway. Proc. Natl. Acad. Sci. U. S. A. 2007, 104 (30), 12341–12346.

(19) Arosio, P.; Knowles, T. P. J.; Linse, S. On the Lag Phase in Amyloid Fibril Formation. Phys. Chem. Chem. Phys. 2015, 17 (12), 7606–7618.

(20) Cohen, S. I. A.; Arosio, P.; Presto, J.; Kurudenkandy, F. R.; Biverstal, H.; Dolfe, L.; Dunning, C.; Yang, X.; Frohm, B.; Vendruscolo, M.; Johansson, J.; Dobson, C. M.; Fisahn, A.; Knowles, T. P. J.; Linse, S. A Molecular Chaperone Breaks the Catalytic Cycle That Generates Toxic Aβ Oligomers. Nat. Struct. Mol. Biol. 2015, 22 (3), 207–213.

(21) Cohen, S. I. A.; Cukalevski, R.; Michaels, T. C. T.; Šarić, A.; Törnquist, M.; Vendruscolo, M.; Dobson, C. M.; Buell, A. K.; Knowles, T. P. J.; Linse, S. Distinct Thermodynamic Signatures of Oligomer Generation in the Aggregation of the Amyloid-β Peptide. Nat. Chem. 2018, 10 (5), 523–531.

(22) Scheidt, T.; Łapińska, U.; Kumita, J. R.; Whiten, D. R.; Klenerman, D.; Wilson, M. R.; Cohen, S. I. A.; Linse, S.; Vendruscolo, M.; Dobson, C. M.; Knowles, T. P. J.; Arosio, P. Secondary Nucleation and Elongation Occur at Different Sites on Alzheimer’s Amyloid-β Aggregates. Science Advances 2019, 5 (4), eaau3112.

(23) Cao, Y.; Tang, X.; Yuan, M.; Han, W. Computational Studies of Protein Aggregation Mediated by Amyloid: Fibril Elongation and Secondary Nucleation. Prog. Mol. Biol. Transl. Sci. 2020, 170, 461–504.

(24) Törnquist, M.; Michaels, T. C. T.; Sanagavarapu, K.; Yang, X.; Meisl, G.; Cohen, S. I. A.; Knowles, T. P. J.; Linse, S. Secondary Nucleation in Amyloid Formation. Chem. Commun. 2018, 54 (63), 8667–8684.

(25) Wang, Q.; Shah, N.; Zhao, J.; Wang, C.; Zhao, C.; Liu, L.; Li, L.; Zhou, F.; Zheng, J. Structural, Morphological, and Kinetic Studies of β-Amyloid Peptide Aggregation on Self-Assembled Monolayers. Phys. Chem. Chem. Phys. 2011, 13 (33), 15200–15210.

(26) Zhao, J.; Wang, Q.; Liang, G.; Zheng, J. Molecular Dynamics Simulations of Low-Ordered Alzheimer β-Amyloid Oligomers from Dimer to Hexamer on Self-Assembled Monolayers. Langmuir 2011, 27 (24), 14876–14887.

(27) Auer, S.; Trovato, A.; Vendruscolo, M. A Condensation-Ordering Mechanism in Nanoparticle-Catalyzed Peptide Aggregation. PLoS Comput. Biol. 2009, 5 (8), e1000458.

(28) Vácha, R.; Frenkel, D. Relation between Molecular Shape and the Morphology of Self-Assembling Aggregates: A Simulation Study. Biophys. J. 2011, 101 (6), 1432–1439.

(29) Morriss-Andrews, A.; Shea, J.-E. Kinetic Pathways to Peptide Aggregation on Surfaces: The Effects of β-Sheet Propensity and Surface Attraction. J. Chem. Phys. 2012, 136 (6), 065103.

(30) Vácha, R.; Linse, S.; Lund, M. Surface Effects on Aggregation Kinetics of Amyloidogenic Peptides. J. Am. Chem. Soc. 2014, 136 (33), 11776–11782.

(31) Do, T. D.; LaPointe, N. E.; Nelson, R.; Krotee, P.; Hayden, E. Y.; Ulrich, B.; Quan, S.; Feinstein, S. C.; Teplow, D. B.; Eisenberg, D.; Shea, J.-E.; Bowers, M. T. Amyloid β-Protein C-Terminal Fragments: Formation of Cylindrins and β-Barrels. J. Am. Chem. Soc. 2016, 138 (2), 549–557.

(32) Shen, L.; Adachi, T.; Vanden Bout, D.; Zhu, X.-Y. A Mobile Precursor Determines Amyloid-β Peptide Fibril Formation at Interfaces. J. Am. Chem. Soc. 2012, 134 (34), 14172–14178.

(33) Radic, S.; Davis, T. P.; Ke, P. C.; Ding, F. Contrasting Effects of Nanoparticle-Protein Attraction on Amyloid Aggregation. RSC Adv. 2015, 5 (127), 105498.

(34) Shezad, K.; Zhang, K.; Hussain, M.; Dong, H.; He, C.; Gong, X.; Xie, X.; Zhu, J.; Shen, L. Surface Roughness Modulates Diffusion and Fibrillation of Amyloid-β Peptide. Langmuir 2016, 32 (32), 8238–8244.

(35) Lin, Y.-C.; Li, C.; Fakhraai, Z. Kinetics of Surface-Mediated Fibrillization of Amyloid-β (12–28) Peptides. Langmuir 2018, 34 (15), 4665–4672.

(36) Šarić, A.; Buell, A. K.; Meisl, G.; Michaels, T. C. T.; Dobson, C. M.; Linse, S.; Knowles, T. P. J.; Frenkel, D. Physical Determinants of the Self-Replication of Protein Fibrils. Nat. Phys. 2016, 12 (9), 874–880.

(37) Gudowska-Nowak, E.; Oliveira, F. A.; Wio, H. S. Editorial: The Fluctuation-Dissipation Theorem Today. Frontiers in Physics 2022, 10. 10.3389/fphy.2022.859799.

(38) Milo, R.; Phillips, R. Cell Biology by the Numbers; Garland Science, 2015

(39) Hayden, E. Y.; Teplow, D. B. Amyloid β-Protein Oligomers and Alzheimer’s Disease. Alzheimers. Res. Ther. 2013, 5 (6), 60.

(40) Selkoe, D. J.; Hardy, J. The Amyloid Hypothesis of Alzheimer’s Disease at 25 Years. EMBO Mol. Med. 2016, 8 (6), 595–608.

(41) Lu, W.; Bueno, C.; Schafer, N. P.; Moller, J.; Jin, S.; Chen, X.; Chen, M.; Gu, X.; Davtyan, A.; de Pablo, J. J.; Wolynes, P. G. OpenAWSEM with Open3SPN2: A Fast, Flexible, and Accessible Framework for Large-Scale Coarse-Grained Biomolecular Simulations. PLoS Comput. Biol. 2021, 17 (2), e1008308.

(42) Ma, Y.-W.; Lin, T.-Y.; Tsai, M.-Y. Fibril Surface-Dependent Amyloid Precursors Revealed by Coarse-Grained Molecular Dynamics Simulation. Front Mol Biosci 2021, 8, 719320.

(43) Chen, W.; Lu, W.; Wolynes, P. G.; Komives, E. A. Single-Molecule Conformational Dynamics of a Transcription Factor Reveals a Continuum of Binding Modes Controlling Association and Dissociation. Nucleic Acids Res. 2021, 49 (19), 11211–11223.

(44) Jin, S.; Bueno, C.; Lu, W.; Wang, Q.; Chen, M.; Chen, X.; Wolynes, P. G.; Gao, Y. Computationally Exploring the Mechanism of Bacteriophage T7 gp4 Helicase Translocating along ssDNA. Proc. Natl. Acad. Sci. U. S. A. 2022, 119 (32), e2202239119.

(45) Chen, X.; Lu, W.; Tsai, M.-Y.; Jin, S.; Wolynes, P. G. Exploring the Folding Energy Landscapes of Heme Proteins Using a Hybrid AWSEM-Heme Model. J. Biol. Phys. 2022. 10.1007/s10867-021-09596-3.

(46) Zheng, W.; Tsai, M.-Y.; Chen, M.; Wolynes, P. G. Exploring the Aggregation Free Energy Landscape of the Amyloid-β Protein (1-40). Proc. Natl. Acad. Sci. U. S. A. 2016, 113 (42), 11835–11840.

(47) Zheng, W.; Tsai, M.-Y.; Wolynes, P. G. Comparing the Aggregation Free Energy Landscapes of Amyloid Beta(1-42) and Amyloid Beta(1-40). J. Am. Chem. Soc. 2017, 139 (46), 16666–16676.

(48) Eastman, P.; Swails, J.; Chodera, J. D.; McGibbon, R. T.; Zhao, Y.; Beauchamp, K. A.; Wang, L.-P.; Simmonett, A. C.; Harrigan, M. P.; Stern, C. D.; Wiewiora, R. P.; Brooks, B. R.; Pande, V. S. OpenMM 7: Rapid Development of High Performance Algorithms for Molecular Dynamics. PLOS Computational Biology. 2017, p e1005659. 10.1371/journal.pcbi.1005659.

(49) Davtyan, A.; Schafer, N. P.; Zheng, W.; Clementi, C.; Wolynes, P. G.; Papoian, G. A. AWSEM-MD: Protein Structure Prediction Using Coarse-Grained Physical Potentials and Bioinformatically Based Local Structure Biasing. J. Phys. Chem. B 2012, 116 (29), 8494–8503.

(50) Tsai, M.-Y.; Zheng, W.; Balamurugan, D.; Schafer, N. P.; Kim, B. L.; Cheung, M. S.; Wolynes, P. G. Electrostatics, Structure Prediction, and the Energy Landscapes for Protein Folding and Binding. Protein Sci. 2016, 25 (1), 255–269.

(51) Sirovetz, B. J.; Schafer, N. P.; Wolynes, P. G. Protein Structure Prediction: Making AWSEM AWSEM-ER by Adding Evolutionary Restraints. Proteins 2017, 85 (11), 2127–2142.

(52) Chen, M.; Lin, X.; Lu, W.; Schafer, N. P.; Onuchic, J. N.; Wolynes, P. G. Template-Guided Protein Structure Prediction and Refinement Using Optimized Folding Landscape Force Fields. J. Chem. Theory Comput. 2018, 14 (11), 6102–6116.

(53) Tsai, M.-Y.; Zhang, B.; Zheng, W.; Wolynes, P. G. Molecular Mechanism of Facilitated Dissociation of Fis Protein from DNA. J. Am. Chem. Soc. 2016, 138 (41), 13497–13500.

(54) Tsai, M.-Y.; Zheng, W.; Chen, M.; Wolynes, P. G. Multiple Binding Configurations of Fis Protein Pairs on DNA: Facilitated Dissociation versus Cooperative Dissociation. J. Am. Chem. Soc. 2019, 141 (45), 18113–18126.

(55) Potoyan, D. A.; Zheng, W.; Komives, E. A.; Wolynes, P. G. Molecular Stripping in the NF-κB/IκB/DNA Genetic Regulatory Network. Proc. Natl. Acad. Sci. U. S. A. 2016, 113 (1), 110–115.

(56) Zhang, B.; Zheng, W.; Papoian, G. A.; Wolynes, P. G. Exploring the Free Energy Landscape of Nucleosomes. J. Am. Chem. Soc. 2016, 138 (26), 8126–8133.

(57) Jin, S.; Contessoto, V. G.; Chen, M.; Schafer, N. P.; Lu, W.; Chen, X.; Bueno, C.; Hajitaheri, A.; Sirovetz, B. J.; Davtyan, A.; Papoian, G. A.; Tsai, M.-Y.; Wolynes, P. G. AWSEM-Suite: A Protein Structure Prediction Server Based on Template-Guided, Coevolutionary-Enhanced Optimized Folding Landscapes. Nucleic Acids Res. 2020, 48 (W1), W25–W30.

(58) Smaoui, M. R.; Poitevin, F.; Delarue, M.; Koehl, P.; Orland, H.; Waldispühl, J. Computational Assembly of Polymorphic Amyloid Fibrils Reveals Stable Aggregates. Biophys. J. 2013, 104 (3), 683–693.

(59) Cecchini, M.; Rao, F.; Seeber, M.; Caflisch, A. Replica Exchange Molecular Dynamics Simulations of Amyloid Peptide Aggregation. J. Chem. Phys. 2004, 121 (21), 10748–10756.

(60) Schwierz, N.; Frost, C. V.; Geissler, P. L.; Zacharias, M. From Aβ Filament to Fibril: Molecular Mechanism of Surface-Activated Secondary Nucleation from All-Atom MD Simulations. J. Phys. Chem. B 2017, 121 (4), 671–682.

(61) Lin, T.-Y.; Ma, Y.-W.; Tsai, M.-Y. Early-Stage Oligomerization of Prion-like Polypeptides Reveals the Molecular Mechanism of Amyloid-Disrupting Capacity by Proline Residues. J. Phys. Chem. B 2023, 127 (5), 1074–1088.

(62) Charest, N.; Tro, M.; Bowers, M. T.; Shea, J.-E. Latent Models of Molecular Dynamics Data: Automatic Order Parameter Generation for Peptide Fibrillization. J. Phys. Chem. B 2020, 124 (37), 8012–8022.

(63) Scherer, M. K.; Trendelkamp-Schroer, B.; Paul, F.; Pérez-Hernández, G.; Hoffmann, M.; Plattner, N.; Wehmeyer, C.; Prinz, J.-H.; Noé, F. PyEMMA 2: A Software Package for Estimation, Validation, and Analysis of Markov Models. J. Chem. Theory Comput. 2015, 11 (11), 5525–5542.

(64) Kumar, S.; Rosenberg, J. M.; Bouzida, D.; Swendsen, R. H.; Kollman, P. A. THE Weighted Histogram Analysis Method for Free-Energy Calculations on Biomolecules. I. The Method. Journal of Computational Chemistry. 1992, pp 1011–1021. 10.1002/jcc.540130812.

(65) Thacker, D.; Barghouth, M.; Bless, M.; Zhang, E.; Linse, S. Direct Observation of Secondary Nucleation along the Fibril Surface of the Amyloid β 42 Peptide. Proc. Natl. Acad. Sci. U. S. A. 2023, 120 (25), e2220664120.

(66) Jean, L.; Lee, C. F.; Vaux, D. J. Enrichment of Amyloidogenesis at an Air-Water Interface. Biophys. J. 2012, 102 (5), 1154–1162.

(67) Schwierz, N.; Frost, C. V.; Geissler, P. L.; Zacharias, M. Dynamics of Seeded Aβ40-Fibril Growth from Atomistic Molecular Dynamics Simulations: Kinetic Trapping and Reduced Water Mobility in the Locking Step. J. Am. Chem. Soc. 2016, 138 (2), 527–539.

(68) Watanabe-Nakayama, T.; Sahoo, B. R.; Ramamoorthy, A.; Ono, K. High-Speed Atomic Force Microscopy Reveals the Structural Dynamics of the Amyloid-β and Amylin Aggregation Pathways. Int. J. Mol. Sci. 2020, 21 (12). 10.3390/ijms21124287.

(69) Zhang, R.; Jalali, S.; Dias, C. L.; Haataja, M. P. Growth Kinetics of Amyloid-like Fibrils: An Integrated Atomistic Simulation and Continuum Theory Approach. PNAS Nexus 2024, 3 (2), gae045.

(70) Bouchard, M.; Zurdo, J.; Nettleton, E. J.; Dobson, C. M.; Robinson, C. V. Formation of Insulin Amyloid Fibrils Followed by FTIR Simultaneously with CD and Electron Microscopy. Protein Sci. 2000, 9 (10), 1960–1967.

(71) Kurouski, D.; Dukor, R. K.; Lu, X.; Nafie, L. A.; Lednev, I. K. Normal and Reversed Supramolecular Chirality of Insulin Fibrils Probed by Vibrational Circular Dichroism at the Protofilament Level of Fibril Structure. Biophys. J. 2012, 103 (3), 522–531.

(72) Usov, I.; Adamcik, J.; Mezzenga, R. Polymorphism Complexity and Handedness Inversion in Serum Albumin Amyloid Fibrils. ACS Nano 2013, 7 (12), 10465–10474.

(73) Xu, Y.; Knapp, K.; Le, K. N.; Schafer, N. P.; Safari, M. S.; Davtyan, A.; Wolynes, P. G.; Vekilov, P. G. Frustrated Peptide Chains at the Fibril Tip Control the Kinetics of Growth of Amyloid-β Fibrils. Proceedings of the National Academy of Sciences 2021, 118 (38), e2110995118.

(74) Zimmermann, M. R.; Bera, S. C.; Meisl, G.; Dasadhikari, S.; Ghosh, S.; Linse, S.; Garai, K.; Knowles, T. P. J. Mechanism of Secondary Nucleation at the Single Fibril Level from Direct Observations of Aβ42 Aggregation. J. Am. Chem. Soc. 2021, 143 (40), 16621–16629.

(75) Okumura, H.; Itoh, S. G. Structural and Fluctuational Difference between Two Ends of Aβ Amyloid Fibril: MD Simulations Predict Only One End Has Open Conformations. Sci. Rep. 2016, 6, 38422.

(76) Heldt, C. L.; Zhang, S.; Belfort, G. Asymmetric Amyloid Fibril Elongation: A New Perspective on a Symmetric World. Proteins 2011, 79 (1), 92–98.

