## Supporting information for "Exploring Abeta42 Monomer Diffusion Dynamics on Fibril Surfaces through Molecular Simulations"

[†]*Contrituted equally to this work*

**Supporting Information** 1

Exploring Abeta42 Monomer Diffusion Dynamics on Fibril Surfaces through Molecular Simulations 1

Fibril structure modeling 2

Figure S1. Example structure of long protofibril 2

Figure S2. The distribution of sampling windows for Abeta monomer diffusion simulation on fibril surfaces. 3

Figure S3. Trajectory-based Nematic Order Parameter (P2) of C-ter Fibril Surface 4

Figure S4. Trajectory-based Nematic Order Parameter of N-ter Fibril Surface 5

Figure S5. Trajectory-based MSD of C-ter Fibril Surface 6

Figure S6. Segment-based MSD of monomers in parallel orientations on the C-terminal fibril surface. 7

Figure S7. Segment-based MSD of monomers in perpendicular orientations on the C-terminal fibril surface. 8

Figure S8. Trajectory-based MSD of N-ter Fibril Surface 9

Figure S9. Trajectory-based mean squared displacement (MSD) of Twisted Fibril 10

Numerical Fitting of Free Energy Profiles from Figure 4 11

### Fibril structure modeling

We used CreateFibril.py, which is a tool to build atomic resolution models of protein fibrils. To initiate the simulation, the input filename (PDB file) is specified using the "-f" parameter (e.g., python CreateFibril.py -f 2MXU.pdb). Following this, the "-c" parameter is employed to select the chains to be replicated and extended; for instance, -c 2,3 designates chains B and C for replication. Subsequently, the "-r" parameter dictates the number of replications for the chosen chains (e.g., -r 30 replicates chains B and C 30 times). The rotation angle of monomers during aggregation is controlled by the "-a" parameter, with -a 0 indicating no rotation. To define the structural characteristics, the total number of residues (amino acids) in the protein is set using the "-nr" parameter (e.g., -nr 32 for a protein with 32 residues). The coordinates and residue point where the fibril axis passes are specified through the "-fp" parameter (e.g., -fp -0.062,-7.195,14.462,26).

Executing the complete command integrates all specified parameters (e.g., python CreateFibril.py -f 2MXU.pdb -c 2,3 -r 30 -a 0 -fp -0.062,-7.195,14.462,26 -nr 32), resulting in the generation of a simulated amyloid fibril with extended protein chains, controlled rotation angles, and defined fibril axis coordinates.

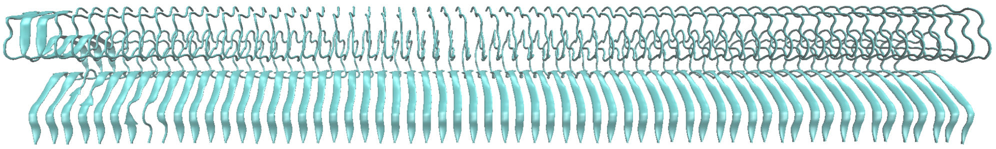
(a)

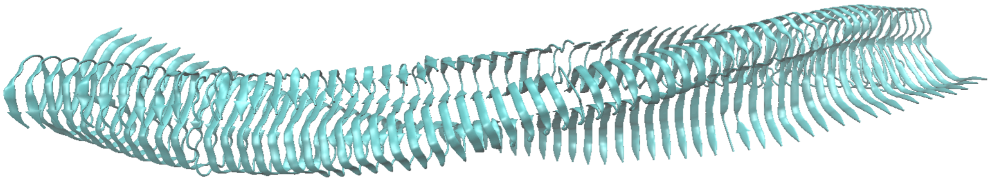
(b)

**Figure S1. Example structure of long protofibril**

(a) Straight fibril
(b) Twist fibril

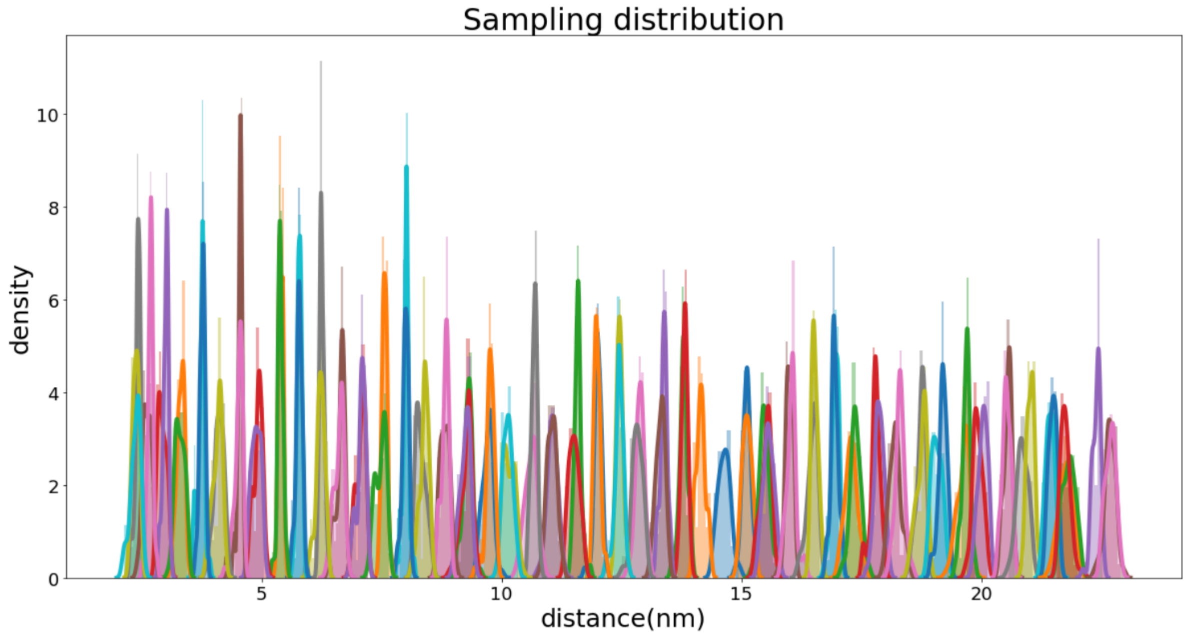


**Figure S2. The distribution of sampling windows for Abeta monomer diffusion simulation on fibril surfaces.**

The Abeta monomer (unbiased) is positioned at different distances from the fibril's centroid along the surface of the fibril. The data shown ensures that the free energy calculation is converged.

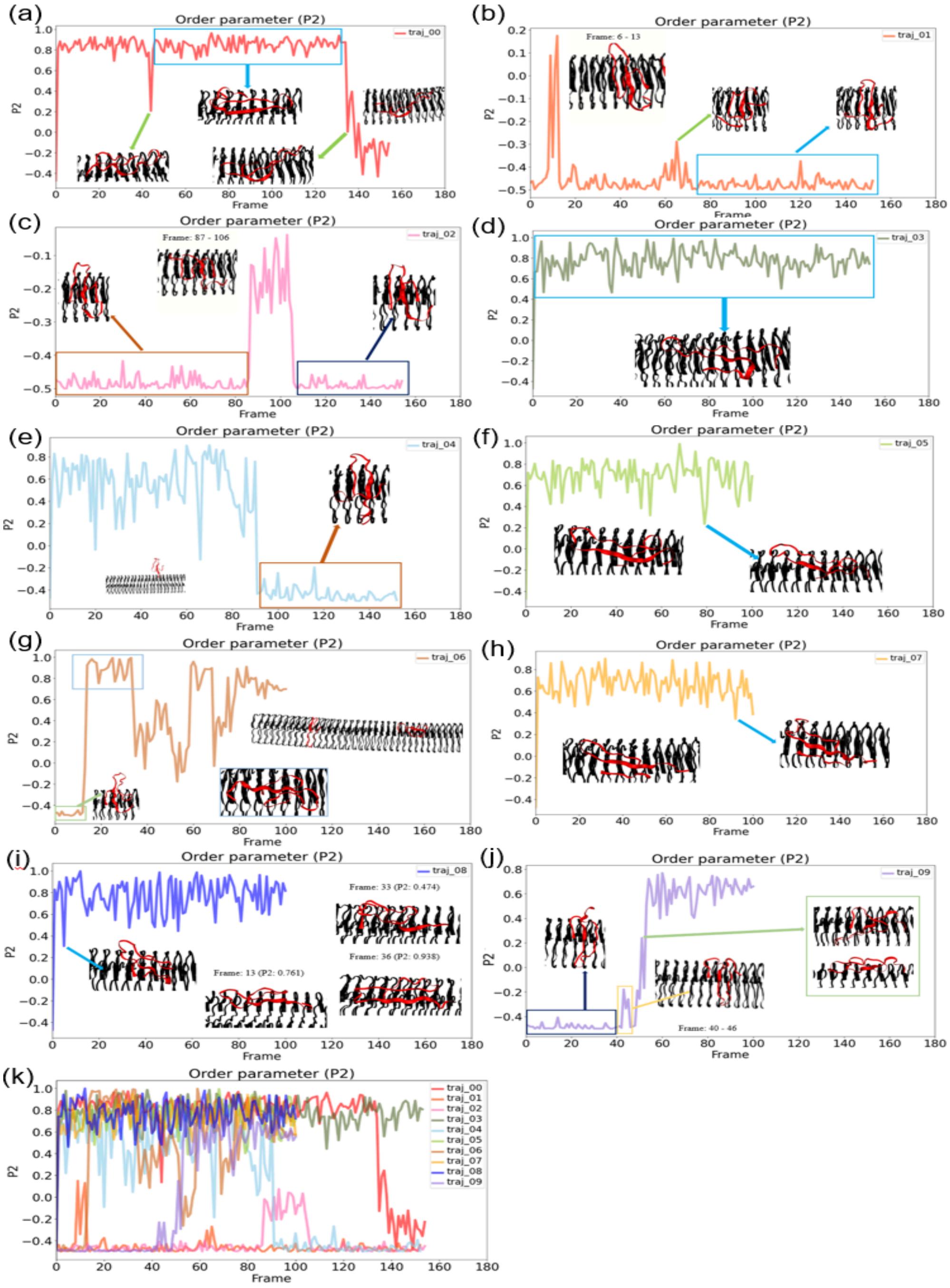

**Figure S3. Trajectory-based Nematic Order Parameter (P2) of C-ter Fibril Surface**

The simulation places monomers above the C-terminal fiber surface. The nematic order parameters (P2) were calculated for each trajectory: (a)-(j) Each panel represents the P2 values of different trajectories over time, with the trajectories labeled from 0 to 9. (See Table 1) (k) Nematic order parameter (P2) of all 10 trajectories are shown together

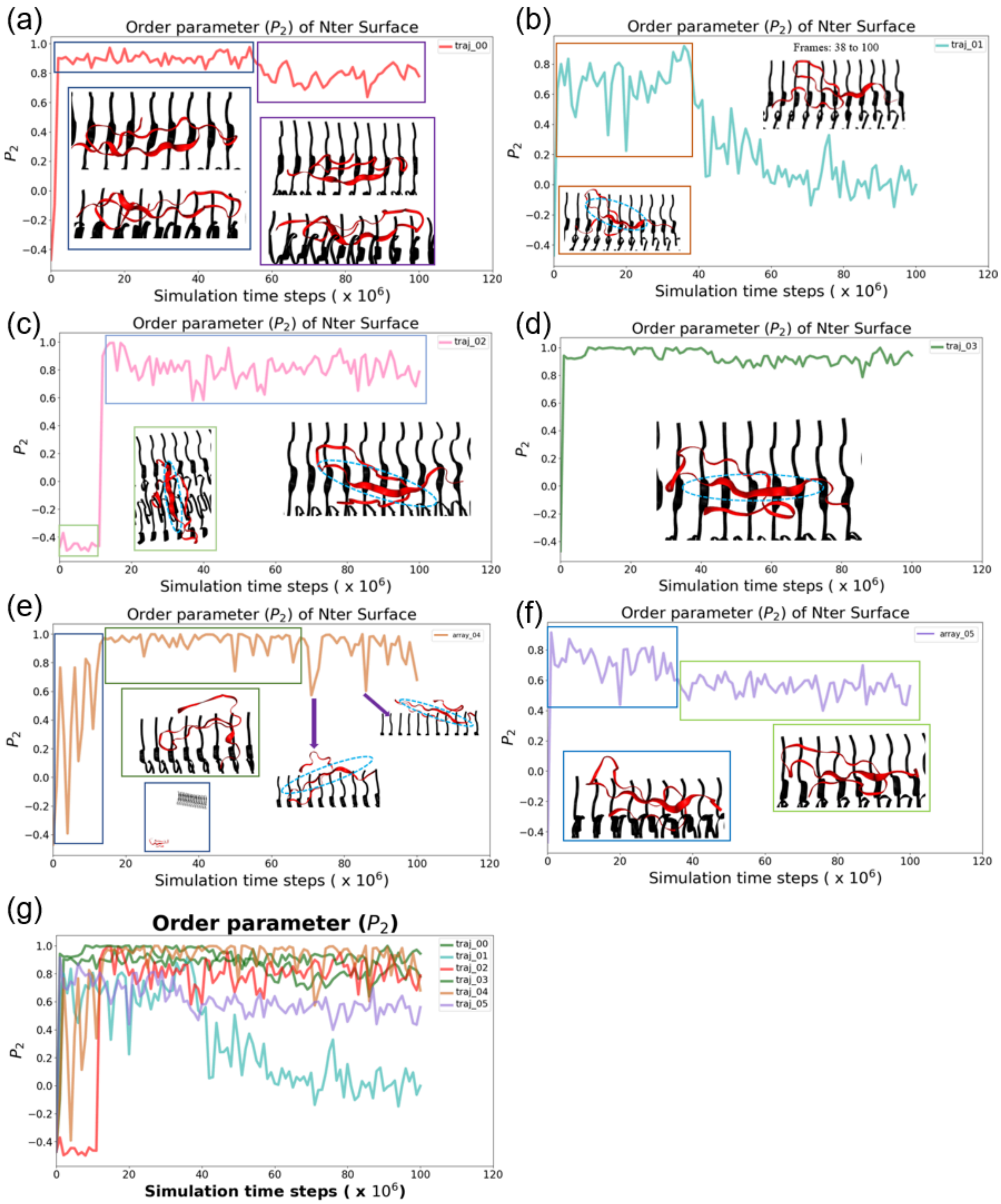


**Figure S4. Trajectory-based Nematic Order Parameter of N-ter Fibril Surface**

The simulation places monomers above the N-terminal fiber surface. The nematic order parameters (P2) were calculated for each trajectory: (a)-(f) Each panel represents the P2 value of different trajectories over time, with the trajectories labeled from 0 to 5. (g) Nematic order parameter of all 6 trajectories are shown together

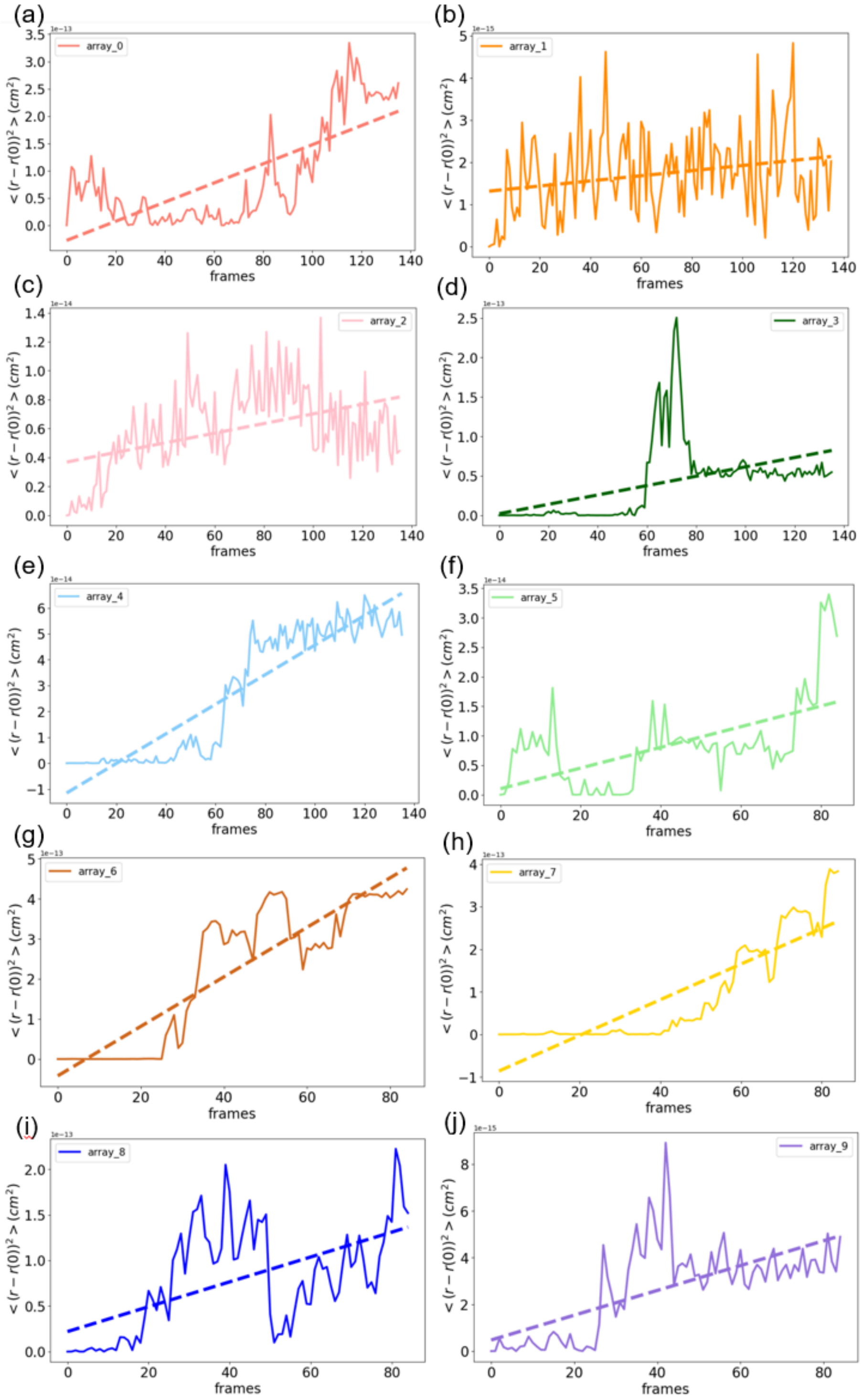


**Figure S5. Trajectory-based MSD of C-ter Fibril Surface**

The simulation places monomers above the C-terminal fiber surface. The MSD was calculated for each trajectory: (a)-(j) Each panel represents the MSD of different trajectories over time, with the trajectories labeled from 0 to 9. (See Table 1)

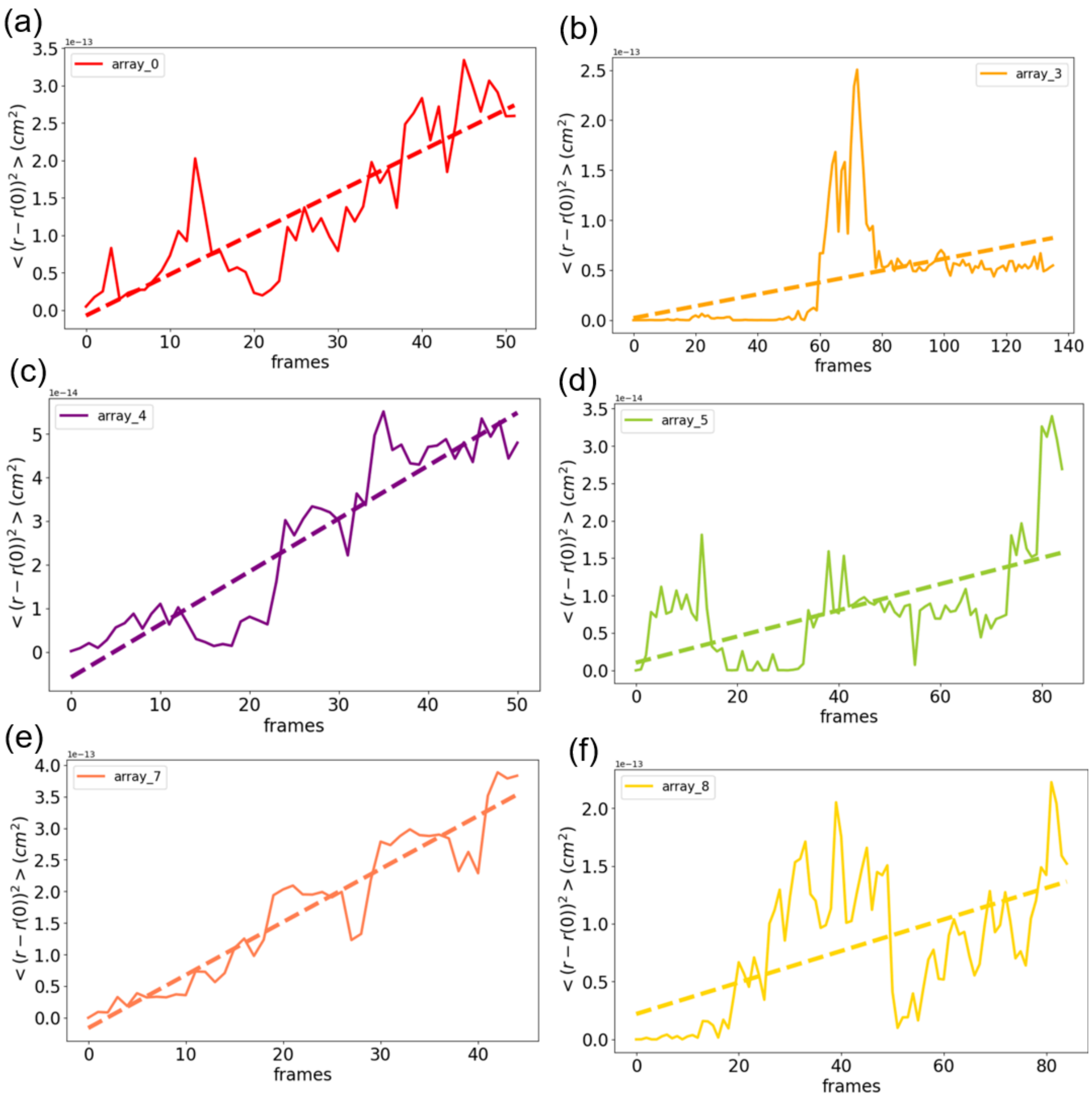


**Figure S6. Segment-based MSD of monomers in parallel orientations on the C-terminal fibril surface.**

The simulation places monomers above the C-terminal fiber surface, selecting trajectories where the monomer orientation is parallel to the main axis of the fiber. The MSD was calculated for each trajectory segment. Each panel shows the MSD of different trajectories over time, with the trajectories labeled from 0 to 9 (See Table 1). The diffusion coefficients (D) calculated for each panel are as follows: (a) 55.1 $\mu m^2/s$ (b) 5.93 $\mu m^2/s$ (c) 12.1 $\mu m^2/s$ (d) 1.75 $\mu m^2/s$ (e) 84.0 $\mu m^2/s$ (f) 13.6 $\mu m^2/s$.

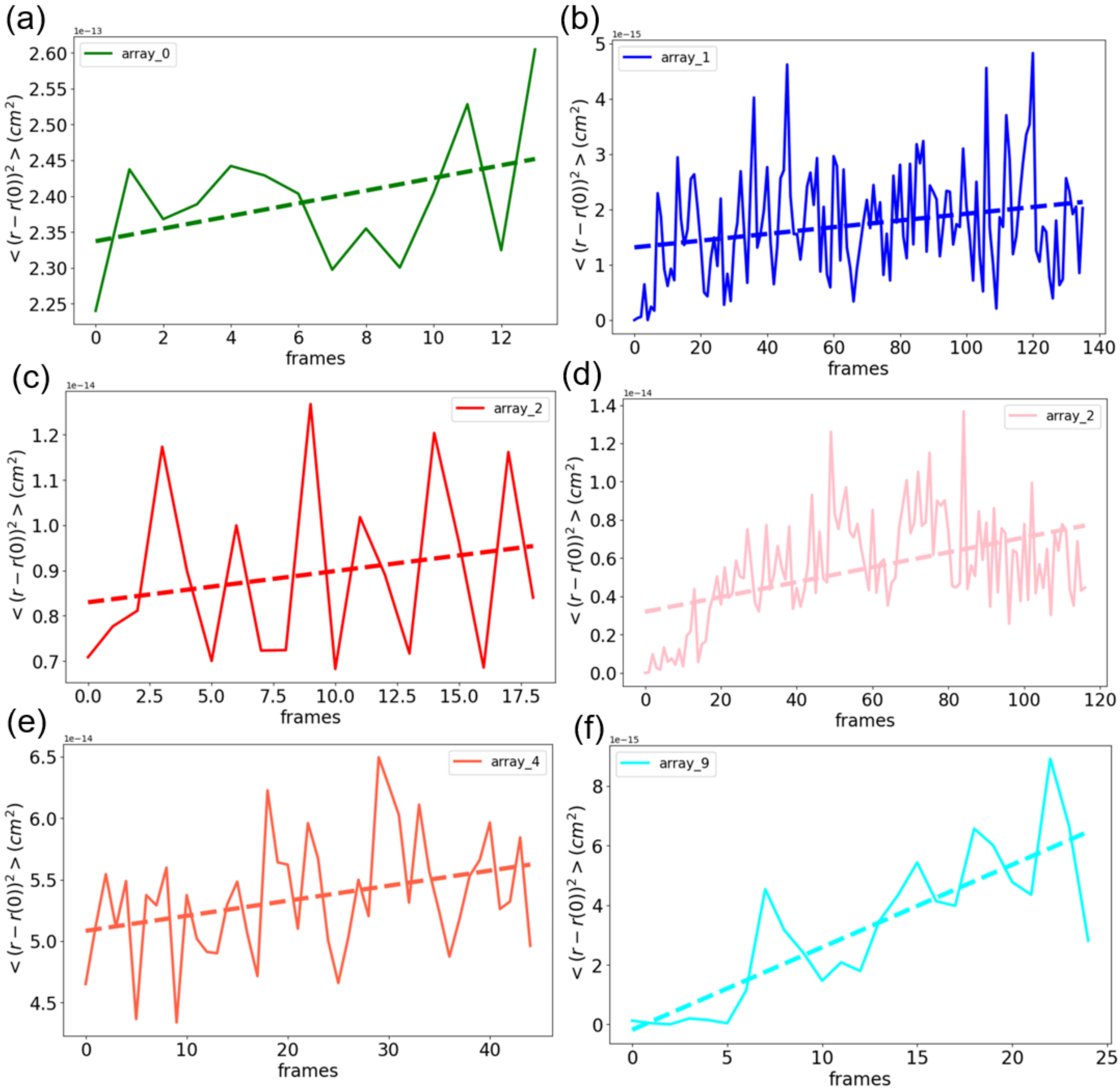


**Figure S7. Segment-based MSD of monomers in perpendicular orientations on the C-terminal fibril surface.**

The simulation places monomers above the C-terminal fiber surface, selecting trajectories where the monomer orientation is perpendicular to the main axis of the fiber. The MSD was calculated for each trajectory segment. Each panel shows the MSD of different trajectories over time, with the trajectories labeled from 0 to 9 (See Table 1). The diffusion coefficients (D) calculated for each panel are as follows: (a) 8.83 $\mu m^2/s$ (b) 6.08 x $10^{-2}$ $\mu m^2/s$ (c) 6.90 x $10^{-1}$ $\mu m^2/s$ (d) 3.89 x $10^{-1}$ $\mu m^2/s$ (e) 1.23 x $10^{-1}$ $\mu m^2/s$ (f) 2.77 x $10^{-1}$ $\mu m^2/s$.

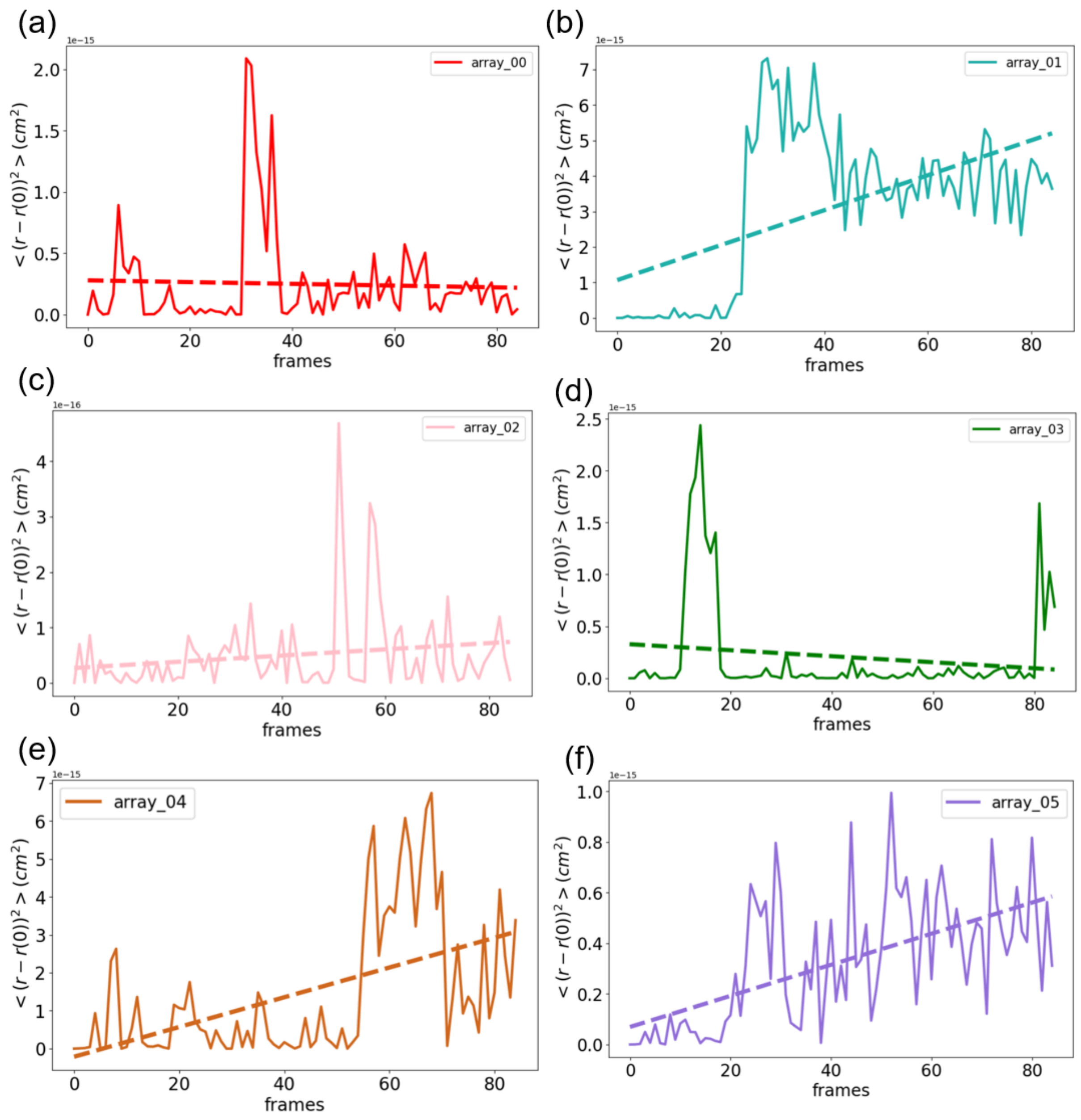


**Figure S8. Trajectory-based MSD of N-ter Fibril Surface**

The simulation places monomers on the N-terminal fiber surface. The MSD was calculated for each trajectory: (a)-(f) Each panel represents the MSD of different trajectories over time, with the trajectories labeled from 0 to 5.

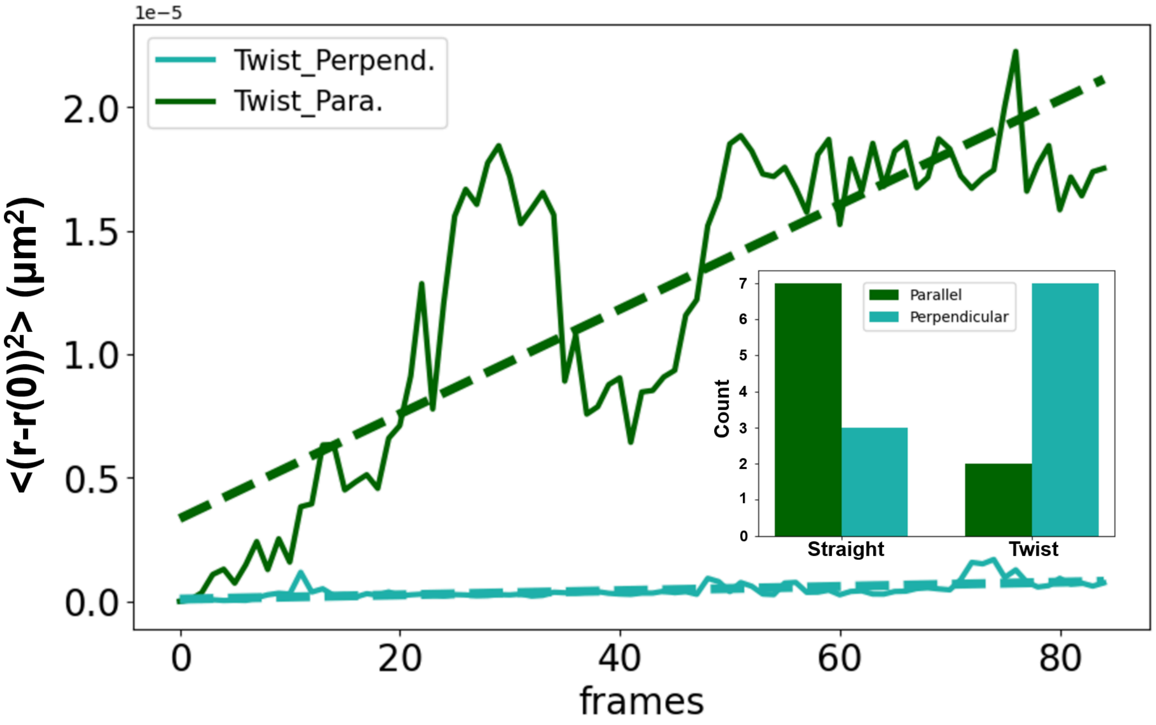


**Figure S9. Trajectory-based mean squared displacement (MSD) of Twisted Fibril**

When monomers are placed above straight and twisted fibril surfaces for 10 simulations each, 7 trajectories exhibit parallel alignment with the straight fibril, while 3 trajectories exhibit perpendicular alignment. Additionally, the panels display the mean squared displacement (MSD) over time, calculated as the average of 2 trajectories aligned parallel to the twisted fibril and 7 trajectories aligned perpendicularly to the fibril. The diffusion coefficients (D) for the parallel and perpendicular are 21.2 $\mu m^2/s$ and 8.58 $\times 10^{-1}$ $\mu m^2/s$.

#### Numerical Fitting of Free Energy Profiles from Figure 4

For the "parallel" monomer, the fitting curve was modeled using the equation:

$$A + B \cdot x + C \cdot cos\,(\omega 1 \cdot x + \phi 1)\,. \qquad \text{(S1)}$$

where the determined fit parameters were: A = -5.06, B = 0.21, C = 0.64, $\omega 1$ = 0.63, $\phi 1$ = 0.42.

For the "perpendicular" monomer, the fitting curve utilized the equation:

$$A + B \cdot x + C \cdot cos\,(\omega 1 \cdot x + \phi 1) + D \cdot sin\,(\omega 2 \cdot x + \phi 2). \qquad \text{(S2)}$$

where the determined fit parameters were: A = -4.36, B = 0.14, C = -0.53, $\omega 1$ = 0.75, $\phi 1$ = -0.23, D = 0.68, $\omega 2$ = -0.37, $\phi 2$ = 1.43.
